# A mammalian-specific domain of MSH5 drives the transition from crossover licensing to designation during meiotic prophase I

**DOI:** 10.64898/2026.06.16.731784

**Authors:** Ky’ara D. Carr, Eliza O’Donnell, Tegan S. Horan, Maame Serwaa Adjei-Boadu, Anna J. Wood, Yongwei Zhang, Winfried Edelmann, Maria de las Mercedes Carro, Paula E. Cohen

**Author notes:** **Corresponding authors**: Maria de las Mercedes Carro and Paula E. Cohen (Lead Contact,).

## Abstract

Meiotic recombination initiates with DNA double-strand breaks (DSBs) repaired as either crossovers (COs) or non-crossovers. Across eukaryotes, MSH4/MSH5 (MutSγ) licenses DSB repair intermediates, directing repair into the class I CO pathway via recruitment of MLH1/MLH3 (MutLγ). In mammals, excess MutSγ sites relative to final MutLγ foci suggest additional MutSγ functions, including directing repair through the minor class II CO pathway. We investigated the role of a mammalian-specific 38-amino acid C-terminal domain of MSH5 using mice lacking this domain (*Msh5^ΔC/ΔC^*). Spermatocytes and oocytes load MSH4 normally to achieve CO licensing in zygonema, but these numbers decline precipitously in pachynema, leading to dramatically reduced MutLγ foci and associated pro-CO factors HEI10 and CNTD1. Despite this, licensing factors RNF212B and MutSγ-associated kinase CDK4 remain persistently upregulated in pachynema. Strikingly, the switch from licensing-associated CDK4 to CO-site-associated CDK2 fails to occur in *Msh5^ΔC/ΔC^* mice, even at residual class I CO events. The result is rapid germ cell death prior to prophase I completion in both sexes. Thus, the loss of the MSH5 C-terminus functionally uncouples the regulatory proteins that define the stepwise patterning of class I COs. Our findings reveal novel early roles for the C-terminus of mammalian MSH5 in converting licensed DSB repair intermediates to designated class I COs.

## INTRODUCTION

Meiosis is the specialized cell division program that generates haploid gametes from diploid progenitor cells, reducing chromosome number by half through two successive divisions — meiosis I and meiosis II — preceded by a single round of DNA replication ^1^. The first division is reductive, separating homologous chromosome pairs, while the second resembles mitosis in separating sister chromatids. The defining stage of meiosis is prophase I, which is subdivided into five stages: leptonema, zygonema, pachynema, diplonema, and diakinesis. During prophase I, homologous chromosomes must find one another, pair, and synapse along their lengths via the formation of a proteinaceous scaffold, the synaptonemal complex (SC) ^2–4^. Indeed, it is the assembly of the SC that provides the cytogenetic roadmap to delineate the unique stages of prophase I. Concurrent with SC formation is the induction of programmed DNA double-strand breaks (DSBs) which form in early leptonema through the action of SPO11 and its partner proteins ^5–9^. Synapsis then facilitates the repair of these DSBs through homologous recombination, ultimately generating crossovers (COs) that provide the physical linkages (chiasmata) necessary for accurate homolog segregation at meiosis I ^1^. Errors in prophase I, and particularly in crossing over, are a leading cause of aneuploidy and, consequently, infertility, miscarriage, and chromosomal disorders such as Trisomy 21 ^10,11^.

Studies over the past three decades have revealed a critical role for meiosis-specific components of the DNA mismatch repair (MMR) family in DSB repair and crossing over across diverse species. In mammals, multiple heterodimers of MutS homologs (MSH) and of MutL homologs (MLH/PMS) exist to coordinate the varied functions of the MMR pathway, from the simple repair of post-replicative errors to the suppression of recombination between similar but non-identical DNA sequences, to their critical roles in meiotic recombination ^12^. In the context of somatic cell repair, MSH2-MSH6 (MutSα) and MSH2-MSH3 (MutSβ) heterodimers interact with two distinct heterodimers of MLH, MLH1-PMS1 (MutLα), MLH1-MLH2 (MutLβ), to achieve these functions. However, during meiosis, a unique subset of heterodimers are employed, and these are MSH4-MSH5 (MutSγ) and MLH1-MLH3 (MutLγ) ^13–16^. The importance of MutSγ and MutLγ for meiosis is underscored by the conserved function of one or both heterodimers in meiotic prophase I in budding yeast, plants, nematodes, and mammals. First identified in budding yeast as being critical for meiosis ^17^, MSH4 and MSH5 have no specific mismatch repair activity despite being highly related to other MutS homologs ^18,28^. Of the MutLγ constituents, on the other hand, MLH1 is shared across all MutL heterodimers, while MLH3 appears to be largely restricted to meiosis in yeast, plant, mouse, and human ^16,18–21^. Importantly, all MutS and MutL orthologs are highly conserved, with MutL homologs sharing 30-40% identity between yeast and human and MutS homologs sharing 20-30% identity across the same evolutionary distance, with the notable exception of a mammalian-specific 38-amino acid C-terminal extension in MSH5 that is absent in all non-mammalian species examined.

DSB repair during meiosis results in the formation of noncrossovers (NCOs) or COs arising through the sequential action of multiple DNA repair pathways ^22,23^. Interestingly, in the mouse, though 250-300 DSBs form in early leptonema, only around 10% of them will become COs, the remainder being processed as NCOs. How this selection for crossing over is achieved is poorly understood, as is the mechanisms by which crossovers conform to the rules of crossover homeostasis that ensure that each chromosome receives at least one CO and that COs are appropriately spaced from each other ^1,24,25^. In *M. musculus*, *A. thaliana*, and *S. cerevisiae*, the majority of COs are generated through the class I pathway involving the MutLγ endonuclease complex ^13,16,18,21,26–30^. MutLγ is recruited to the site of the DSB repair by MutSγ, and canonically these two complexes are presumed to act together at all times, consistent with the stoichiometry and mode of action of all MutS and MutL heterodimers ^15,31–37^. Indeed, in budding yeast, MSH4 and MSH5 were considered founding members of the ZMM network of pro-crossover factors that are specifically critical for class I CO formation ^38–40^. However, studies from our lab and others have revealed unexpected roles for mammalian MutSγ beyond its recruitment of MutLγ. Most prominently, during prophase I in male mice, MSH4 and MSH5 accumulate in discrete foci along chromosome axes in zygonema, with a maximal frequency of around 150 foci per nucleus^35,36,41,42^. By contrast, MLH1 and MLH3 appear on chromosome axes considerably later, at pachynema, and their focus frequency matches the final tally of class I CO sites at 23-25 foci in mouse spermatocytes ^21,37,43–45^. Thus, while the final deposition of MutLγ marks the ultimate “designation” of class I CO from the larger pool of DSB sites, the accumulation of MutSγ at these sites earlier in prophase I is considered to represent “licensing” of certain DSB repair intermediates towards the class I CO pathway. However, in the context of mammalian meiosis, these licensed sites can still be repaired through the utilization of other pathways, including mitosis-defined structure specific nuclease resolution pathway, known as the class II CO pathway, driven by MUS81-EME1 and representing fewer than 5-10% of all COs in mouse ^46^.

In concordance with the temporal differences in accumulation of MutSγ and MutLγ, the phenotypic loss of MSH4/MSH5 is more drastic in mice than that of MLH1/MLH3, though in all cases the resulting homozygous mutants are infertile ^21,36,41,45,47^. Thus, mice harboring deletions in MutSγ components show meiotic disruption in zygonema, characterized by failure to complete synapsis, persistent unrepaired DSBs (decorated with the RecA homolog, RAD51), and wide-scale apoptosis of spermatocytes before pachynema in adult males, and of oocytes at around the point of dictyate arrest (the stage at which fetal oocytes arrest in prophase I at the time of birth) ^35,36,41,42^. By contrast, loss of either MutLγ components allows full pachytene progression, complete synapsis, and removal of RAD51, but a failure to establish class I COs, leading to a reduction in chiasmata by 90-95% by diplonema ^21,27,47–49^. Taken together, these observations lead us to hypothesize a role for MutSγ beyond its role in recruiting MutLγ to nascent class I CO sites, and instead suggest that mammalian MutSγ may participate in additional DSB repair events that can give rise to COs and NCOs through other repair pathways. In line with this, previous work from our lab showed that a point mutant variant of *Msh5* in the mouse that lacks a functional ATPase domain allows for progression of prophase I through until diakinesis, but results in the loss of all COs regardless of their route of generation from the DSB precursor ^42^.

To further explore the mammalian-specific functions of MutSγ, we turned to the 38-amino acid sequence in the C-terminal region of the MSH5 protein that is highly conserved amongst mammals, including non-eutherian mammals, but is not present in other species. We generated a mutant mouse model lacking this C-terminal region of *Msh5* (designated *Msh5*^ΔC^). Analysis of homozygous mutant *Msh5*^ΔC/ΔC^ animals reveals that MutSγ accumulates normally in zygonema in males and females along with the critical MutSγ-associated CO licensing factor, RNF212B. However, the C-terminal region of MSH5 is essential for continued association of MutSγ with DSB repair intermediates in pachynema, and for processing of DSB repair intermediates towards crossing over. As a result, MutSγ foci decline dramatically in pachynema, leading to reduced pachytene recruitment of MutLγ and its associated pro-CO factors, HEI10 and CNTD1 ^50–53^. Interestingly, RNF212B ^54,55^ and the MutSγ-associated kinase, CDK4 ^56^, remain elevated through pachynema, despite the reduction in MutSγ foci, essentially disconnecting MutSγ from its associated licensing factors. At the same time, despite persistence of almost half the normal class I designated CO sites, marked with MLH1, MLH3, CNTD1, and HEI10, we observe no loading of the CO-site associated kinase, CDK2 ^56^, in pachynema. Thus, we propose that the transition from CO licensing to CO designation is associated with a critical switch in kinase regulation from CDK4 to CDK2, and that this switch is dependent on the full function of MSH5, including activities involving the C-terminus of the mammalian protein. As a result, *Msh5*^ΔC/ΔC^ male and female meiocytes display meiotic failure, leading to loss of post-meiotic germ cells in the testis, and a dramatic depletion of the ovarian reserve at birth. Additionally, our work supports the idea that the complete function of MutSγ is required for the resolution of all DSB intermediates from pachynema onwards, including those assigned to class I and class II CO pathways.

## RESULTS

### MutSγ localization pattern is normal in zygonema, but it is compromised in *Msh5*^ΔC/ΔC^ spermatocytes during pachynema

To further investigate the role of MutSγ in mouse meiosis, we utilized a novel mouse mutant line bearing a truncated C-terminal region of MSH5 (*Msh5*^ΔC^) (**Fig. S1**). This truncation removes a highly conserved 38 amino acid sequence present in mammals (**Fig. S1C**). To examine MutSγ localization in spermatocytes from homozygous mutant animals (*Msh5*^ΔC/ΔC^), we used antibodies against MSH4, the obligate partner of MSH5 ^35,36,42^, on chromosome prophase I spread preparations (**Fig. 1A-J**). To stage the spermatocytes, we used co-immunofluorescence (IF) staining against the lateral/axial element of the SC, SYCP3 (Synaptonemal complex protein 3). In spermatocytes from adult *Msh5^+/+^* male mice, MSH4 localizes to the chromosome cores of the SC starting in zygonema ^35^. The number of MutSγ foci peaks at the end of this stage-beginning of pachynema with an average foci number of 125.4 ± 13.20 foci per nucleus (**Fig. 1B, K**) and then declines to an average of 61.06 ± 33.54 foci per nucleus during total pachynema (**Fig. 1C-D, L**). There was no significant change in total MSH4 foci per nucleus between wild-type and *Msh5*^ΔC/ΔC^ mice at zygonema (**Fig. 1A-B, F-G, K**). Analysis of prophase I in *Msh5*^ΔC/ΔC^ mice revealed that pachytene spermatocytes show a large number of unpaired (“asynapsed”) chromosomes; therefore, cells displaying at least 10 fully synapsed chromosomes were categorized as “pachytene-like” (**Fig. 1H-I**), while cells displaying bulging ends, desynapsis, and/or short length of the pseudoautosomal region (PAR) were defined as “diplotene-like” (**Figure 1J**). Moreover, by pachynema, MSH4 focus counts were significantly reduced in *Msh5*^ΔC/ΔC^ spermatocytes compared to their *Msh5*^+/+^ counterparts by approximately 64% (**Fig. 1L**). In both *Msh5*^+/+^ and *Msh5*^ΔC/ΔC^ mice, MSH4 foci were absent from diplotene and diplotene-like spermatocytes (**Fig. 1E, J**). These observations confirm that the *Msh5*^ΔC/ΔC^ mouse line generates a truncated version of MSH5 capable of forming a heterodimer with MSH4 and localizing to the chromosome axes in zygonema at normal frequencies, but the heterodimer is unable to remain associated with the chromosome axes in pachynema. More specifically, the data suggest that the C-terminal domain of MSH5 is critical to promote stable MutSγ association with CO sites, permitting the distinction between the role of MSH5 in CO designation at pachynema from its earlier function in crossover licensing during early zygonema.

**Figure 1.**
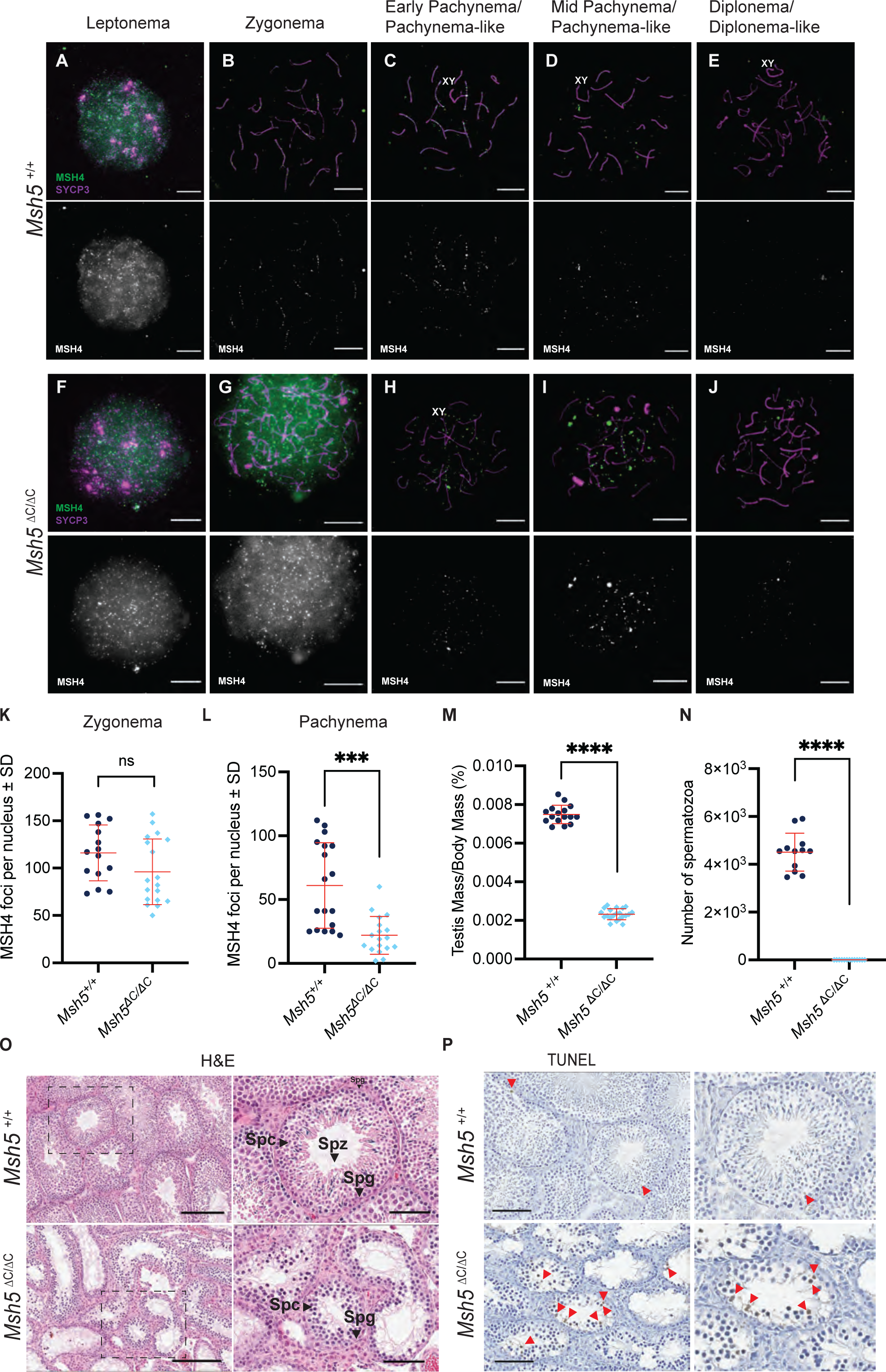
*Msh5 ^ΔC/ΔC^* meiocytes exhibit a significant decrease in MSH4 foci. (A-J) Representative images of spermatocyte prophase I spreads from *Msh5^+/+^*and *Msh5^ΔC/ΔC^* mice immunostained against MSH4 (green) and SYCP3 (magenta, chromosome cores). White scale bars are equal to 10μm. (K-L) Quantification of MSH4 foci per nucleus in zygotene (K) and pachytene or pachytene-like stage (L). Each point represents the average of foci per nucleus, and bars represent average foci number ± SD, n = 18 cells from 3 mice. Statistical comparisons were performed using the Mann-Whitney test; p-values are either shown as p < 0.001 (***) or ns = not significant. (M) Testis weights relative to body mass for *Msh5^+/+^* and *Msh5^ΔC/ΔC^* male mice. Dots represent data points for individual mice and bars represent the average foci ± SD, n = 16 and n = 21, respectively. (N) Cauda epididymal spermatozoa counts obtained by swim out from *Msh5^+/+^* and *Msh5^ΔC/ΔC^* mice, n = 18 cells from 3 mice. Dots represent data points for individual mice, and bars represent the average foci ± SD, n = 16 and n = 21, respectively. Data were analyzed by the Mann-Whitney test (****p <0.0001). (O) Representative histological sections of testis from *Msh5^+/+^*and *Msh5^ΔC/ΔC^* mice stained with Hematoxylin-Eosin, bars indicate 200 μm. Spc: Spermatocyte, Spz: Spermatozoa, Spg: Spermatogonia. (P) Representative images of TUNEL staining for each genotype, bars indicate 200 μm. The insert shows magnification of the *Msh5^ΔC/ΔC^* panel; red arrowheads indicate apoptotic cells.

### Truncation of the C-terminal domain of MSH5 leads to altered gonadal architecture and germ cell loss in male mice

To assess the impact of the C-terminal truncation of MSH5 on gametogenesis, we assessed the reproductive phenotype of *Msh5*^+/+^ and *Msh5*^ΔC/ΔC^ mice. Compared to wild-type littermates, *Msh5*^ΔC/ΔC^ male mice showed a reduction in testis weights by almost 70% and the complete absence of cauda epididymal spermatozoa **(Fig. 1M, N),** similar to what we previously observed for *Msh5^-/-^* animals ^36^. Seminiferous tubules of *Msh5* ^ΔC/ΔC^ males exhibited a loss of germ cells from the prophase I layer of cells onwards, with no visible spermatozoa (**Fig. 1O**). As expected, no pups were born from crosses between *Msh5*^ΔC/ΔC^ males with wild-type or *Msh5*^ΔC/ΔC^ females (**Fig. S2**). To assess cell death by apoptosis, we performed TUNEL staining of paraffin-embedded testis sections, revealing a drastic increase in cellular death in the prophase I layer of the seminiferous epithelium of *Msh5*^ΔC/ΔC^ mice compared to wild-type littermates (**Fig. 1P, red arrowheads**). Taken together, these data demonstrate that *Msh5*^ΔC/ΔC^ male mice show spermatogenic failure in prophase I, leading to the complete absence of post-meiotic haploid germ cells.

### DSBs form normally in *Msh5* ^ΔC/ΔC^ spermatocytes, but fail to repair in a timely fashion

To examine DSB induction and initial processing in *Msh5*^ΔC/ΔC^ males, we first monitored DSB induction through staining with an antibody against the phosphorylated form of histone H2AX, known as γH2AX, which marks the position of unrepaired DSBs ^57–59^, along with SYCP3. γH2AX is widely distributed and detectable throughout the nucleus during leptonema in wild-type spermatocytes. As prophase I progresses, DSBs become progressively processed, leading to a gradual reduction in γH2AX signal (**Fig. S3A-D**). By pachynema, all DSBs are now repaired, resulting in the almost complete loss of γH2AX signal on the autosomes and persistence on the XY-containing sex body by pachytene and diplotene stages (**Fig. S3C-D**). By contrast, pachytene-like spermatoctyes from *Msh5*^ΔC/ΔC^ spermatocytes show initially normal distribution of γH2AX signal from leptonema onwards, but persistent autosomal γH2AX signal from pachytene-like cells, continuing into the diplotene-like stage, indicating either the persistence of unrepaired DSBs into late prophase I, or the formation of additional DSBs after the first round of DSB induction in leptonema (**Fig. S3E-H**).

To explore early DSB repair events in *Msh5*^ΔC/ΔC^ spermatocytes, we stained meiotic spreads using antibodies against RAD51 and SYCP3. RAD51 is a recombinase enzyme that is recruited to single-stranded DNA along with its meiosis-specific homolog DMC1 to catalyze homology search and strand exchange, which is essential for the early stages of DSB repair upstream of different NCO and CO pathways ^60,61^. In wild-type spermatocytes, RAD51 appears as individual foci along chromosome axes soon after DSB induction, being found at maximal frequency between leptonema to zygonema (**Fig. 2A-B, I**). As DSB repair proceeds, the number of RAD51 diminishes to almost zero in pachynema and diplonema, with any residual foci being largely restricted to the sex chromosomes (**Fig. 2C-D, J**).

**Figure 2.**
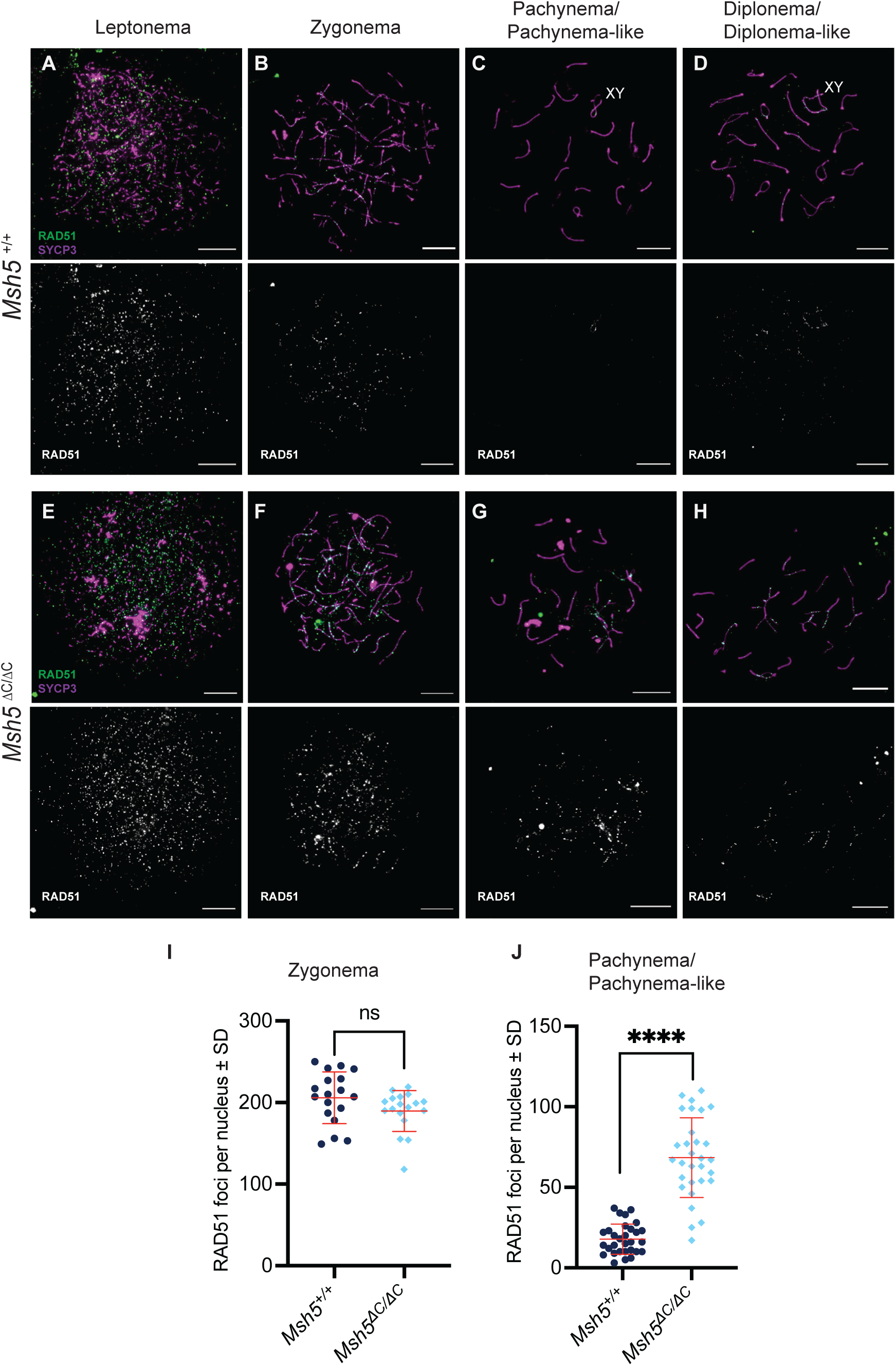
DSBs remain unrepaired in pachytene-like *Msh5^ΔC/ΔC^* spermatocytes. (A-H) Representative images of spermatocyte prophase I spreads from *Msh5^+/+^*and *Msh5^ΔC/ΔC^* mice immunostained against RAD51 (green) as a proxy for DSB repair and SYCP3 (magenta). White scale bars are equal to 10μm. (I-J) Quantification of total RAD51 foci per zygotene nucleus (I) and pachytene and pachytene-like nucleus (J). Each point represents the average of foci per nucleus, and bars represent average foci number ± SD, n = 18 cells from 3 mice for *Msh5^+/+^* and n = 30 cells from 3 mice for *Msh5^ΔC/ΔC^*, respectively. Statistical comparisons were performed using the Mann-Whitney test; p-values are either shown as p < 0.0001 (****) or ns = not significant.

In line with the normal induction of DSBs, we observed normal numbers of RAD51 in zygotene spermatocytes from *Msh5* ^ΔC/ΔC^ mice (**Fig. 2E-F, I)**. However, in pachytene-like and diplotene-like cells from *Msh5*^ΔC/ΔC^ males, RAD51 focus frequency was significantly elevated above that of wild-type male mice (**Fig. 2G-H, J)**. Taken together, our data suggest that early DSB formation is normal in *Msh5*^ΔC/ΔC^ males, leading to normal accumulation of RAD51 in zygonema. However, we observe a persistence of unrepaired breaks, or the induction of new DSBs labeled with RAD51 in pachynema-like spermatocytes.

### *Msh5*^ΔC/ΔC^ spermatocytes display synapsis defects and increased non-homologous chromosome interactions

We next evaluated the status of the synapsis and SC formation through prophase I, using co-staining of SYCP3 with SYCP1 on meiotic chromosome spreads collected from *Msh5*^+/+^ and *Msh5*^ΔC/ΔC^ animals. SYCP1, a component of the transverse filaments of the SC, acts as a zipper-like filament linking homologous chromosomes together during meiosis ^62^. In wild-type mice, SYCP3 is distributed across the nucleus during leptonema as discrete threads associating with compacting chromosomes (**Fig. S4A**). As cells progress into zygonema, continuous synapsis between homologous chromosomes while SYCP1 threads coalesce into short, and then longer, filaments (**Fig. S4B**). By pachynema, spermatocytes show 19 fully synapsed homologous autosomes and a sex chromosome pair that shows synapsis only at the PAR, with increased desynapsis through until diplonema (**Fig. S4C**). By contrast, spermatocytes from *Msh5*^ΔC/ΔC^ males begin chromosome synapsis and SC formation normally in leptonema and zygonema, but by pachynema they exhibit extensive asynapsis, with some chromosomes able to synapse along their entire lengths and others showing partial pairing and mis-pairing (**Fig S4D-F**).. This aberrant synapsis was also demonstrated through the prescence of HORMAD1 at mis-synapsed regions through until late pachynema. HORMAD1 is an essential SC component that is present throughout most of prophase I in males – it is removed from autosomes by pachynema but is retained on the unpaired X and Y arms. Then, in diplonema, HORMAD1 returns on the desynapsing axes of all chromosomes (**Fig. S4G-H**) ^63^.

### Establishment of class I crossovers is impaired in *Msh5* ^ΔC/ΔC^ spermatocytes

To determine whether the loss of the MSH5 C-terminus disrupts class I CO designation in males, we assessed the localization pattern and frequency of MLH1 and MLH3 foci by IF on SYCP3-stained pachytene spreads prepared from *Msh5*^+/+^ and *Msh5*^ΔC/ΔC^ animals. In wild-type spermatocytes, MLH1 and MLH3 form discrete foci along the SC during pachytene, with average focus counts of 23.11 ± 1.71 and 23.41 ± 2.2, respectively (**Fig. 3A, C, D, F**). In *Msh5*^ΔC/ΔC^ spermatocytes, MLH1 and MLH3 foci numbers are significantly reduced in both the abundance and brightness compared to those found in pachytene spermatocytes from *Msh5*^+/+^ littermates, at 9.73 ± 3.6 and 8.74 ± 4.37, respectively (**Fig. 3B, C, E, F)**. Unexpectedly, this finding differs from our previous observation in *Msh5*^GA/GA^ spermatocytes, which lacks the ATPase activity of MSH5, in which we see a complete absence of MutLγ foci, while *Msh4*^-/-^ and *Msh5*^-/-^ null mutants fail to achieve pachynema and thus also do not load MutLγ ^35,36,42^.

**Figure 3.**
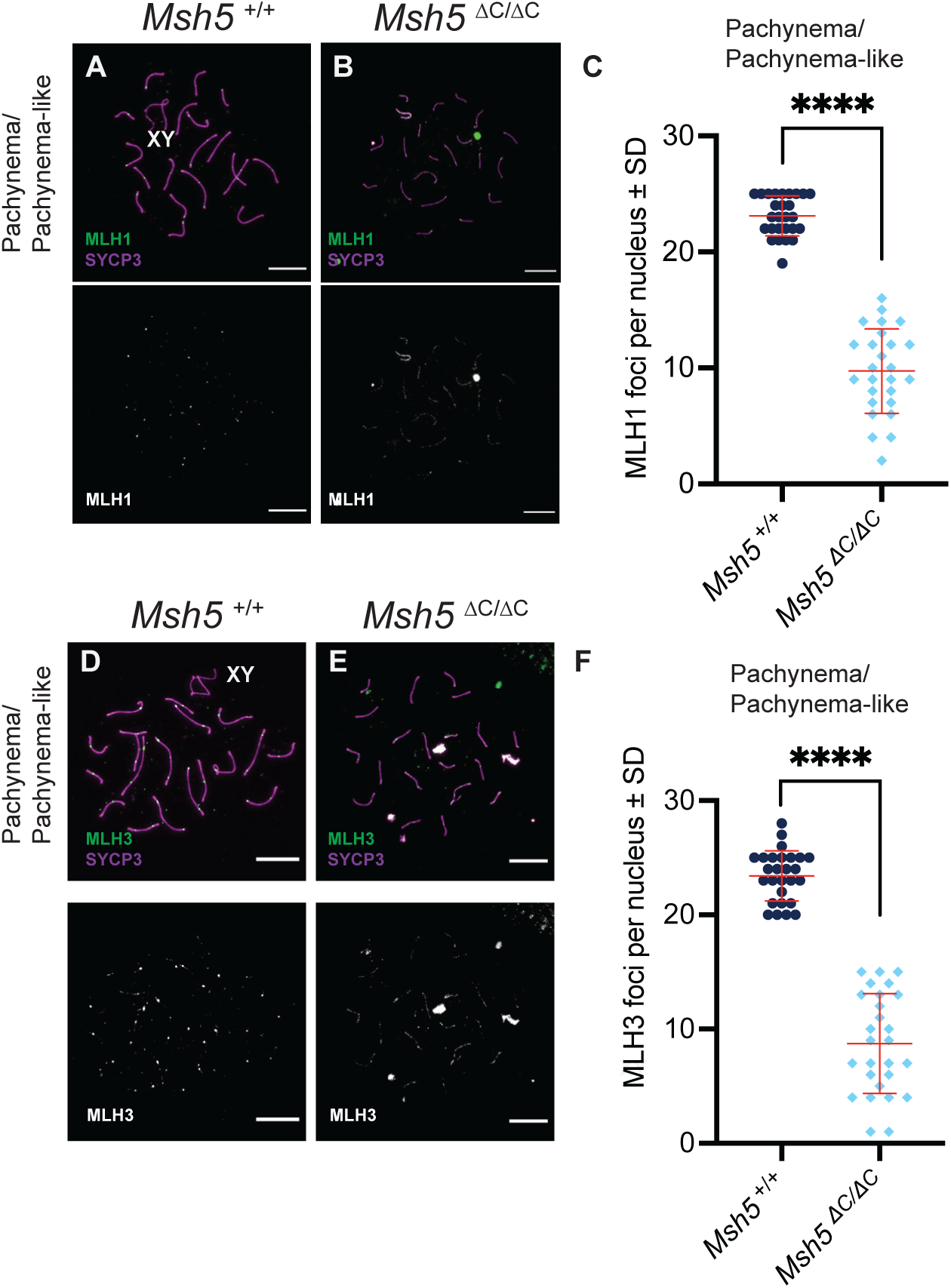
Dynamics of crossover levels differ in *Msh5^ΔC/ΔC^* spermatocytes. (A-B) Representative images of pachytene and pachytene-like spermatocyte prophase I spreads from *Msh5^+/+^* and *Msh5^ΔC/ΔC^* male mice immunostained against MLH1 (green, class I COs) and SYCP3 (magenta). White scale bars are equal to 10 μm. (C) Quantification of MLH1 foci per nucleus. Each point represents the average of foci per nucleus, and bars represent the average foci number ± SD, n = 27 cells from 3 mice. (D-E) Representative images of pachytene and pachytene-like spermatocyte prophase I spreads immunostained against MLH3 (green, class I COs) and SYCP3 (magenta). White scale bars are equal to 10 μm. (F) Quantification of MLH3 foci per nucleus. Each point represents the average of foci per nucleus and bars represent average foci number ± SD, n = 27 cells from 3 mice. Statistical comparisons were performed using the Mann-Whitney test; p-values are either shown as p < 0.0001 (****) or ns = not significant.

Taken together, these findings demonstrate that full MSH5 activity is required for robust MutLγ association with chromosome cores, leading to the full complement of nascent class I CO events. Notably, however, the C-terminal truncation of MSH5 does not completely abolish MutLγ recruitment, as MutSγ remains capable of loading onto the SC and recruiting MutLγ in *Msh5*^ΔC/ΔC^ males. Interestingly, however, when we performed air-dried preparations of diakinesis-staged cells to assess final chiasmata numbers, we found no evidence for diakinesis-stage cells in over 100 cells examined from *Msh5*^ΔC/ΔC^ animals, suggesting that the prophase I disruption in these animals leads to cell death before exit from prophase I. This is in contrast to our previous report exploring the ATPase-deficient *Msh5^GA/GA^* mouse line, in which many diakinesis-staged spermatocytes were observed, but all lacked any residual chiasmata ^42^. These observations suggest that the C-terminal region of MSH5 is necessary for maintaining chromosome associations throughout prophase I, and that the low number of residual class I COs found in these mice is insufficient to ensure persistent connections between homologous chromosomes.

### Localization of CO-promoting factors is compromised in *Msh5* ^ΔC**/**ΔC^ spermatocytes

The recruitment of MutSγ and MutLγ to DSB repair intermediates is a multistep process that requires the action of upstream meiotic recombination factors as well as an emerging group of crossover regulators that are specific to class I (reviewed by ^23^). Crossover-specific stabilization of MutSγ in pachynema is mediated in part by crossover RING proteins (CORs), which share a common domain arrangement composed of an N-terminal RING domain, a coiled-coil domain, and an intrinsically unstructured C-terminus ^64^. In mammals, three COR have been identified: Ring Finger Proteins 212 and 212B (RNF212, RNF212B, putative SUMO and Ubiquitin E3 ligases, respectively) and Human enhancer of invasion-10 (HEI10) ^53–55,65,66^. RNF212 and RNF212B exhibit a broad accumulation pattern in spermatocytes that parallels MutSγ localization ^53,54^, whereas HEI10 becomes selectively enriched at class I CO-designated sites ^53^. In addition to these CORs, Cyclin N-terminal domain-containing-1, CNTD1, also appears to regulate the designation of class I CO sites, possibly through facilitating the efficient removal of RNF212/212B and the subsequent recruitment of HEI10 ^50^.

In wild-type spermatocytes, numerous foci of RNF212B accumulate along SCs in zygonema (**Fig. 4A**), and by late pachynema, only CO-designated sites will retain RNF212B (**Fig. 4B, M**). However, in *Msh5* ^ΔC/ΔC^ spermatocytes, there is a significant increase in foci numbers of RNF212B in pachynema-like stages relative to wild-type pachytene spermatocytes (Fig. **4H, M**), suggesting that removal of RNF212B from zygonema to pachynema is impaired in these cells.

**Figure 4.**
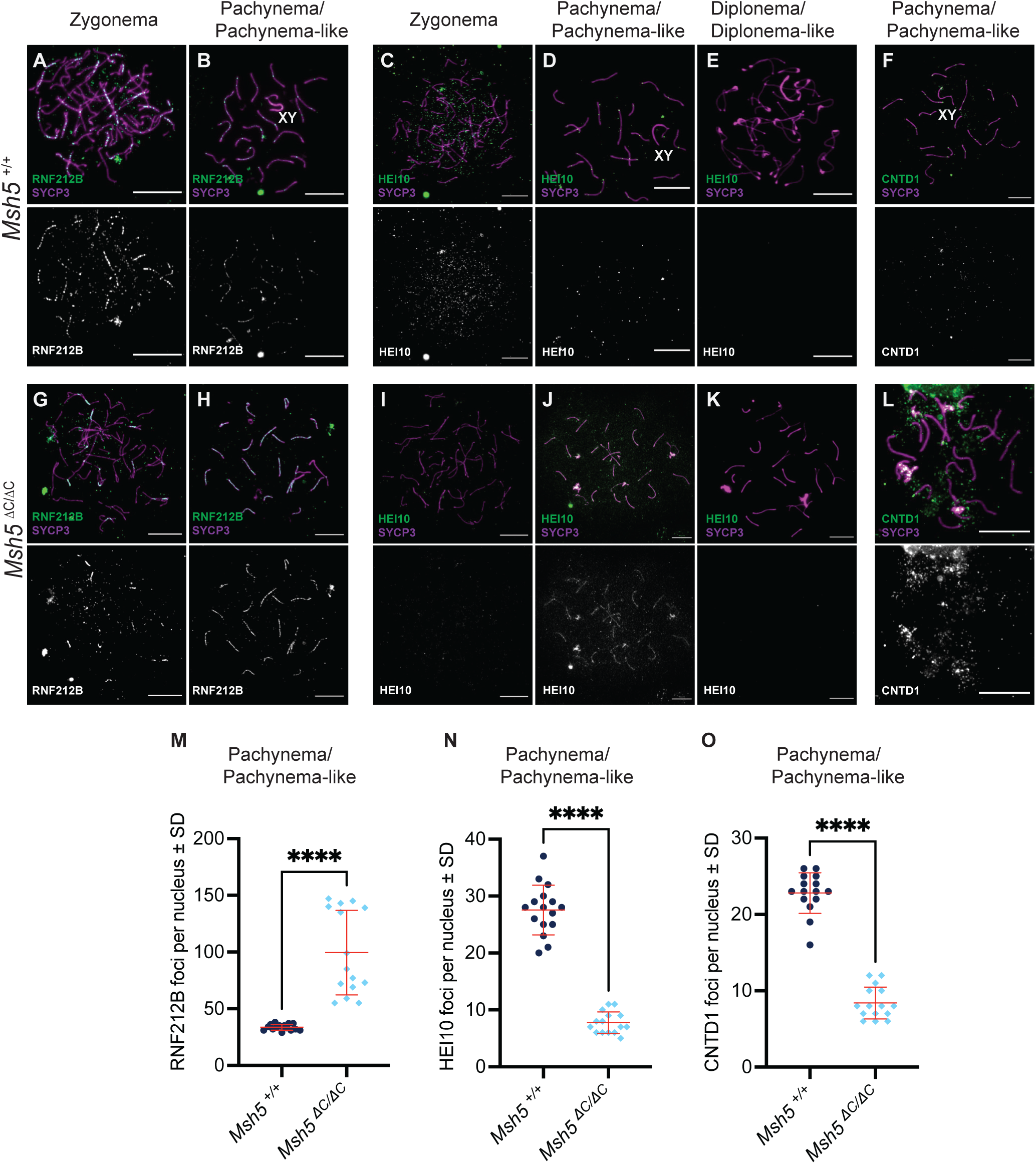
Localization of pro crossover factors is impaired in *Msh5^ΔC/ΔC^* spermatocytes. (A-L) Representative images of pachytene and pachytene-like spermatocyte prophase I spreads from *Msh5^+/+^* and *Msh5^ΔC/ΔC^* male mice immunostained against RNF212B (A-B, G-H), HEI10 (C-E, I-K), and CNTD1 (F, L) in green and SYCP3 in magenta. White scale bars are equal to 10 μm. (E) Quantification (average foci ± SD) of RNF212B foci per nucleus. (M-O) Quantification of foci per nucleus for RNF212B (M) HEI10 (N), and CNTD1 (O). Each point represents the average of foci per nucleus, and bars represent average foci ± SD, n = 15 cells from 3 mice. Statistical comparisons were performed using the Mann-Whitney test; p-values are either shown as p < 0.0001 (****) or ns = not significant.

The switch from CO-licensed to CO-designated sites is marked by a switch in COR protein localization from RNF212/RNF212B to HEI10, which is facilitated by CNTD1 ^50,52^. Therefore, we assessed HEI10 and CNTD1 accumulation during pachynema in *Msh5^+/+^*and *Msh5*^ΔC/ΔC^ spermatocytes (**Fig. 4C-F, I-L, N, O**). As expected, wild-type spermatocytes accumulate HEI10 and CNTD1 foci numbers similar to MLH1 and MLH3 foci numbers by pachynema (27.6 ± 4.3 and 22.8 ± 2.6 foci per nucleus, respectively). Analysis of HEI10 localization in *Msh5*^ΔC/ΔC^ spermatocytes at pachynema reveals a failure to accumulate at CO sites, remaining significantly reduced compared with that in wild-type spermatocytes (7.7 ± 1.9, **Fig. 4I-K, N**). Similarly, CNTD1 localization at CO sites in pachynema was significantly reduced in *Msh5*^ΔC/ΔC^ spermatocytes relative to that observed in wild-type spermatocytes (8.4 ± 2.09, **Fig. 4F, L, O**). Overall, these data indicate that the C-terminal domain of MSH5 is required for stabilization of MutSγ during pachynema and for recruitment of the pro-CO factors CNTD1 and HEI10. As a consequence, disruption of these functions hinders RNF212B removal from the SC and abolishes robust loading of HEI10 and CNTD1 in pachynema.

### *Msh5*^ΔC/ΔC^ spermatocytes exhibit aberrant localization of CDK4 and CDK2

Cyclin-dependent kinases (CDKs) are essential regulators of meiotic recombination, coordinating homolog pairing and synapsis, as well as driving the maturation of recombination intermediates into COs required for accurate chromosome segregation. During prophase I, key CDKs have been identified at distinct stages of synapsis and recombination. In particular, CDK4 is essential for fertility and localizes to synapsed regions of the SC from zygonema to early pachynema, after which it is lost as chromosomes begin to desynapse ^56,67^. It has been hypothesized that CDK4 kinase activity may be required either to stabilize axial-lateral element interactions and promote their association or to destabilize proteins associated with asynapsed axes, facilitating their dissociation upon completion of synapsis. Importantly, CDK4 associates with the SC during zygonema and is lost once CO designation has occurred, exchanged with CDK2 as the sole CO site-associated cyclin-dependent kinase.

In *Msh5^+/+^* male mice, we observe the expected accumulation of CDK4 on chromosome cores from zygonema onwards, initially in high numbers in early pachynema (49.7 ± 17.5) that then decline through to mid-pachynema (6.3 ± 6.1) (**Fig. 5A-B, M**). CDK4 is then absent from late pachynema onwards (**Fig. 5C-D, M**). In *Msh5*^ΔC/ΔC^ spermatocytes, we observed similar zygotene appearance of CDK4, persisting through until early pachynema **(Fig. 5G, M)**, but then continuing to increase in frequency across the chromosome axes in mid-to-late pachytene-like and diplotene-like stages (**Fig. 5H-J, M**), rather than the complete loss of CDK4 from mid-pachynema onwards that was observed in *Msh5^+/+^* spermatocytes at an equivalent stage (**Fig. 5M**).

**Figure 5.**
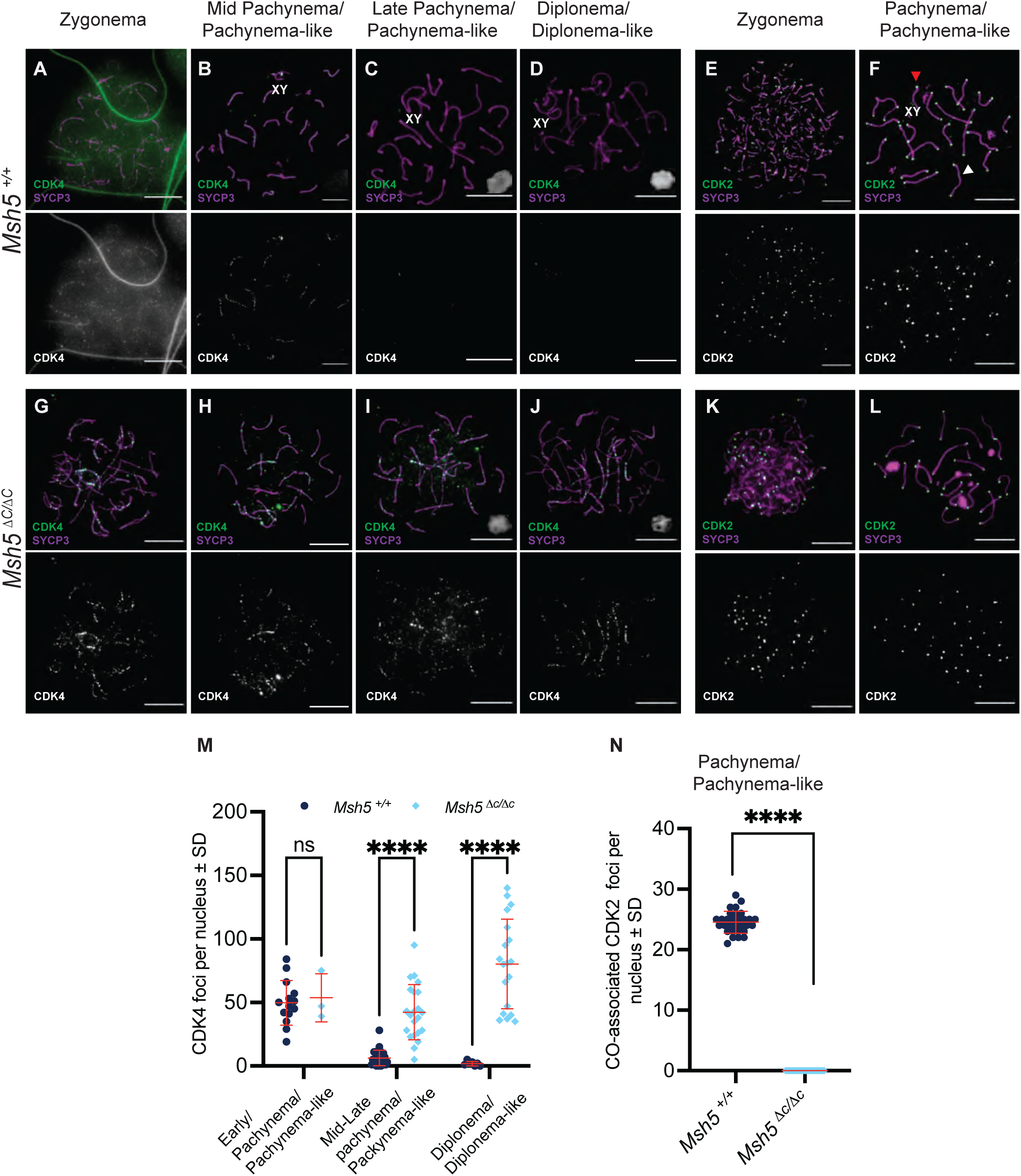
Localization of cyclin-dependent kinases is impaired in *Msh5^ΔC/ΔC^* spermatocytes. (A-L) Representative images of pachytene and pachytene-like prophase I spreads from *Msh5^+/+^* and *Msh5^ΔC/ΔC^* male mice immunostained against CDK4 (A-D, G-J) and CDK2 (E-F, K-L) in green and SYCP3 in magenta. White scale bars are equal to 10 μm. (M) Quantification of foci per nucleus for CDK4 in early and mid-pachynema/pachynema-like cells, and diplonema/diplonema-like cells. Each point represents the average of foci nucleus, and bars represent the average foci number ± SD, n = 17, n = 26, n = 7, respectively, for *Msh5^+/+^* and n = 3, n = 22, n = 19, respectively, for *Msh5^ΔC/ΔC^* from 3 animals. (N) Quantification of foci per nucleus for CDK2 in pachynema/pachynema-like cells. Each point represents the average of the foci nucleus, and bars represent the average foci number ± SD, n = 30 from 3 animals. Statistical comparisons were performed using the Mann-Whitney test; p-values are either shown as p < 0.0001 (****) or ns = not significant.

CDK2 activity is essential for meiotic progression through its localization to telomeres during early prophase I, where it promotes telomere tethering to the nuclear envelope and regulates CO formation at chromosome cores (**Fig. 5E-F, N)** ^68,69^. Additionally, CO site-associated CDK2 localizes to the SC from mid-pachynema to late pachynema, where it is thought to regulate phosphorylation of recombination nodule proteins and promote the recruitment of pro-crossover factors at designated sites ^70^. Importantly, the switch from CDK4 to CDK2 from late zygonema to mid-pachynema is a hallmark feature of the switch from CO licensing to designation. As expected, therefore, we observed the appearance of CDK2 at CO-designated sites from mid-pachymema onwards, in addition to the telomere-associated foci present on the ends of each chromosome in wild type spermatocytes (**Fig. 5E-F**, white and red arrows, respectively). In spermatocytes from *Msh5*^ΔC/ΔC^ mice, while CDK2 localization to telomeric regions is not affected during pachynema, we observed the complete loss of CO site-associated CDK2 (**Fig. 5K-L, N**). Notably, the complete absence of CDK2 despite partial localization of CNTD1 during pachynema in *Msh5*^ΔC/ΔC^ spermatocytes reinforces our earlier assertions that CNTD1 does not act as the CDK2-associated cyclin-like protein in mammalian meiosis in contrast to its ortholog in *C. elegans*, COSA-1, which appears to interact with worm CDK-2 ^50,70^. Collectively, the persistence of CDK4 at synapsed regions of the SC and the loss of CO site-associated CDK2 foci provide insight into the timing of meiotic disruption in these mutants, as CDK4 is normally removed before mid-late pachynema, when CDK2 accumulates in the SCs ^56^ .

### Loss of the MSH5 C-terminal domain leads to female infertility

Compared to male meiosis, there is a lack of understanding of the function of MutSγ during female meiosis. Thus, to address this relative imbalance in our understanding of CO regulation between males and females, we explored the meiotic phenotype of *Msh5*^ΔC/ΔC^ female mice by IF of prophase I oocytes from in embryonic day 18.5 females. First, we observed the localization pattern of MSH4 in oocytes from *Msh5*^ΔC/ΔC^ females compared to wild-type littermates. As previously observed in spermatocytes, oocytes lacking the C-terminal domain of MSH5 can localize MutSγ to the SC during zygonema at similar numbers to wild-type oocytes (117.9 ± 46.95 vs 134.7 ± 40.89, respectively, **Fig. 6A, D, G**). However, the MSH4 foci numbers drastically drop by pachynema in these oocytes, with a 60% reduction compared to wild-type controls (21.33 ± 12.79 compared to 54.33 ± 24.33, **Fig. 6B, E, H**), similar to previously observed for males (**Fig. 1L**). Analysis of markers of synapsis and crossing over revealed a similar phenotype of *Msh5*^ΔC/ΔC^ females to that seen in adult *Msh5*^ΔC/ΔC^ males as well: early DSB induction and repair were normal in *Msh5*^ΔC/ΔC^ females, as demonstrated by normal early accumulation of γH2AX and RAD51 (**Fig. S5A, D, G, J**). However, by pachynema-like stages, we observed increased γH2AX flares across the chromosomes (**Fig. S5B, E, C, F**) along with an increased number of RAD51 foci in pachytene-like and diplotene-like oocytes from *Msh5*^ΔC/ΔC^ females compared to wild-type littermates (**Fig. S5H-I, K-L, M**), indicating that DSB repair is impaired in oocytes with a C-terminal domain-truncated MSH5. When we analyzed localization patterns of SYCP1 and HORMAD1 in *Msh5*^ΔC/ΔC^ oocytes, we found increased accumulation of unsynapsed chromosomes in pachynema-like oocytes compared to those from wild-type littermates (**Fig. S5N-U**), with frequent non-homologous chromosome interactions (**Fig. S5U**, white arrowhead) similar to that observed in *Msh5*^ΔC/ΔC^ males (**Fig. S4H**). As expected from our analysis of CO formation in *Msh5*^ΔC/ΔC^ males, staining using antibodies against the class I CO markers MLH1 and MLH3 showed drastically reduced numbers of foci for both markers in pachytene-like oocytes from *Msh5*^ΔC/ΔC^ females (**Fig. 6I-N**), with means of 26.48 ± 3.15 and 24.94 ± 3.03 respectively for wild-type females and 0.43 ± 0.72 and 0.63 ± 0.85 respectively for *Msh5*^ΔC/ΔC^ females. This decrease is much more pronounced in females than in males, where the MLH1 and MLH3 foci numbers for *Msh5^ΔC/ΔC^* spermatocytes were 9.73 ± 3.6 and 8.74 ± 4.37, respectively.

**Figure 6.**
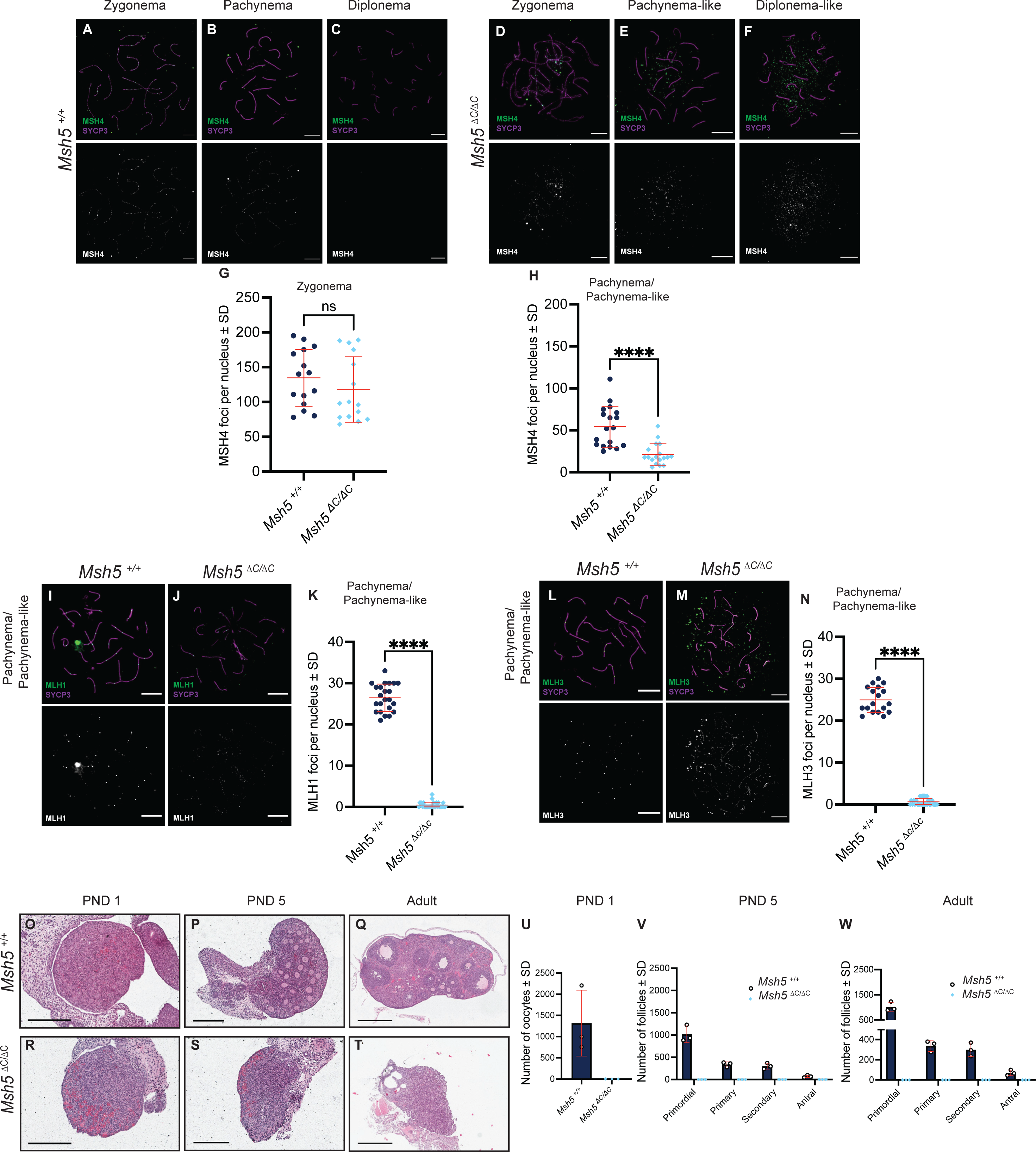
*Msh5^ΔC/ΔC^* females are infertile and recapitulate the spermatocyte prophase I phenotype. (A-F) Representative images of oocyte prophase I spreads from *Msh5^+/+^*and *Msh5^ΔC/ΔC^* 18.5 dpc female embryos immunostained against MSH4 (green) and SYCP3 (magenta). White scale bars are equal to 10μm. (G-H) Quantification of MSH4 foci per nucleus in zygotene (G) and pachytene or pachytene-like stage (H). Each point represents the average of foci per nucleus, and bars represent average foci number ± SD, n = 15 cells from 3 mice. Statistical comparisons were performed using the nonparametric Mann-Whitney test; p-values are either shown as p < 0.0001 (****) or ns = not significant. (I-J) Representative images of oocyte prophase I spreads from *Msh5^+/+^*and *Msh5^ΔC/ΔC^* 18.5 dpc female embryos immunostained against MLH1 (green) and SYCP3 (magenta). White scale bars are equal to 10μm. (K) Quantification of MLH1 foci per nucleus in pachynema and pachynema-like oocytes. Each point represents the average of foci per nucleus, and bars represent the average foci number ± SD, n = 23 cells from 3 mice. (L-M) Representative images of oocyte prophase I spreads from *Msh5^+/+^* and *Msh5^ΔC/ΔC^* 18.5 dpc female embryos immunostained against MLH3 (green) and SYCP3 (magenta). White scale bars are equal to 10μm. (N) Quantification of MLH3 foci per nucleus in pachynema and pachynema-like oocytes. Each point represents the average of foci per nucleus, and bars represent average foci number ± SD, n = 18 cells from 3 mice. Statistical comparisons were performed using the nonparametric Mann-Whitney test; p-values are either shown as p < 0.0001 (****). (O-T) Histological sections of ovaries from *Msh5^+/+^* and *Msh5^ΔC/ΔC^* female mice at post-natal day (PND) 1, 5, and 10 weeks (adult). Scale bars for PND1/PND5 are equal to 200 μm and for adult are equal to 500 μm. (U-W) Quantification of primordial, primary, secondary, and antral follicles in ovaries at PND1 (U), PND5 (V), and 10-week-old female mice (W) from *Msh5^+/+^*and *Msh5^ΔC/ΔC^* lines. Each point represents the total number of oocytes/follicles per ovary, and bars represent the average of oocytes/ follicles ± SD from 3 animals.

We examined ovarian development and folliculogenesis in *Msh5^ΔC/ΔC^* female mice compared to wild-type littermates by quantification of the ovarian follicular composition from postnatal day 1 (PND 1), post-natal day 5 (PND 5), and 10-week-old (adult) females, documenting the follicular stages from primordial (early) follicles through antral (late pre-ovulatory) follicles (**Fig. 6O-W**). At PND1, we observe 1318 ± 777.2 oocytes per ovary in wild-type females (**Fig. 6O, U**), while no oocytes were observed in PND1 ovaries from *Msh5^ΔC/ΔC^* pups (**Fig. 6R, U**). This result was surprising given previous reports from us and others that oocytes persist in *Msh5^-/-^* (and *Msh4^-/-^*) females through at least until week 3 of life ^35,36,41^. This observation remained consistent through early development (PND5) and into adulthood, where we observed no follicles at any stage in the ovaries of *Msh5^ΔC/ΔC^* females, compared to large numbers of follicles at all stages in the ovaries of wild-type littermates (**Fig. 6P-Q, Y, S-T, W**). Consistent with this, we did not obtain pups from crosses between *Msh5^ΔC/ΔC^* females and wild-type males (**Fig. S2**).

To study if the presence of a truncated version of MSH5 disrupts the accumulation of MutSγ-associated licensing and pro-CO factors to the same degree as that reported for male meiosis, above, we studied the localization of RNF212B, CDK4, HEI10, CNTD1, and CDK2 in prophase I chromosome spreads from *Msh5*^ΔC/ΔC^ female embryos and wild-type littermates (**Fig. S6**). Analysis of these markers revealed a similar phenotype to that observed in *Msh5*^ΔC/ΔC^ males for females, including persistence of RNF212B (**Fig. S6A-B, G-H**), and drastically reduced loading of HEI10 and CNTD1 (**Fig. S6C-F, I-L**). Wild-type oocytes accumulate CDK4 on chromosome cores from zygonema onwards, slightly different than spermatocytes. Although foci numbers are initially high in zygonema, it seems that the reduction of CDK4 numbers in oocytes happens later than in spermatocytes, given that late pachynema oocytes still show some levels of CDK4 (**Fig. S6M-O**). Nonetheless, we also observed persistence of CDK4 in diplonema oocytes from *Msh5^ΔC/ΔC^* females (**Fig. S6S-U**). Lastly, we observed a similar pattern of CDK2 localization in wild-type pachytene oocytes compared to spermatocytes with telomeric (**Fig. S6Q**, red arrows) and CO site-associated foci (white arrows). Similar to the phenotype observed in males, *Msh5*^ΔC/ΔC^ pachytene-like oocytes show no change in CDK2 foci in telomeric regions and a complete loss of CO site-associated CDK2 compared to wild-type oocytes (**Fig. S6V)**. Overall, analysis of *Msh5*^ΔC/ΔC^ female mice shows that lost of the C-terminal domain of MSH5 causes a very similar phenotype than males, in which MutSγ can localize to the SC of oocytes in normal numbers by zygonema, however, by pachynema the truncated version of MSH5 is associated with impaired accumulation of pro-CO factors HEI10, CNTD1 and CDK2 at licensed citessites that leads to a drastic loss in MutLγ foci and infertility. Alongside, *Msh5*^ΔC/ΔC^ oocytes show persistence of RNF212B and CDK4 by diplonema, as observed for males, suggesting that the timing for meiotic disruption in females is similar to that of males.

## DISCUSSION

The repair of DNA double-strand breaks (DSBs) during prophase I of meiosis involves a variety of repair machineries that, while conserved across eukaryotes, may be differentially utilized to achieve a fixed and highly regulated distribution of CO events genome-wide that is specific to each species ^23^. In mammals, this CO regulation takes on additional realms of complexity with the well-described sexual dimorphism that exists in the distribution and frequency of CO events across the genome ^71–73^. Moreover, recent studies have also highlighted significant variability in CO regulation across different genetic backgrounds in the mouse ^71,74,75^. In the current study, we investigated the role of the MSH4-MSH5 heterodimer to better define its contribution to these repair decisions in male and female mice.

The conserved function of MutSγ in facilitating meiotic recombination and crossover formation has been demonstrated across diverse species, including *Saccharomyces cerevisiae, Arabidopsis thaliana, Tetrahymena thermophila, Caenorhabditis elegans,* and *Mus musculus* ^15,31,35–37,41,76–79^. Our analysis of MutSγ in mice provides a foundation for understanding the spatial and temporal distribution of MutSγ and its associated factors throughout mammalian meiosis, while also revealing features of its function that might be restricted to mammals. We find that, while homozygous *Msh5*^ΔC*/*Δ^*^C^* meiocytes show normal early DSB repair dynamics and accumulation of MSH4, indicative of normal CO licensing, they exhibit non-homologous chromosome interactions from pachynema onwards, along with persistent RAD51 foci into pachynema, along with aberrant localization of MutLγ and other pro-crossover factors, such as HEI10, and CNTD1. We cannot differentiate between persistent RAD51 indicating the persistence of unrepaired breaks that were formed in leptonema or the formation of *de* novo breaks in zygonema and pachynema. In contrast to the expected 23-25 MutLγ-designated crossover sites present at pachynema, fewer than 12 class I crossover events (around 43% of normal) are detected in pachytene-like spermatocytes from *Msh5*^ΔC*/*Δ^*^C^* mice, with all these markers being found at approximately the same frequency at this stage. Surprisingly, however, three features of pachytene-like cells from *Msh5*^ΔC*/*Δ^*^C^* male mice deviate from both the normal pro-CO dynamics observed in wild-type littermate control males and also other meiotic mutant lines that affect either CO licensing or CO designation steps. First, we observe a persistent elevation of RNF212/RNF212B without a concomitant persistence of MSH4, where usually the two are found together, as in *Cntd1^-/-^* and *Hei10^-/-^* male mice ^54,55^. Second, we observe persistence and upregulation of CDK4 until diplonema. Thirdly, we see a failure to load CDK2 onto any class I-designated sites, even those few sites that persist in *Msh5*^ΔC*/*Δ^*^C^* males. The persistence of RNF212B and CDK4 without MSH4, coupled with the differential localization of pro-CO factors — partial recruitment of HEI10, CNTD1, and MutLγ versus the complete absence of CDK2 — reveals a potential bifurcation in the transition from CO licensing to CO designation. These findings suggest that although loss of the MSH5 C-terminal domain allows limited class I crossover designation, it disrupts a critical meiotic transition from CDK4- to CDK2-dependent regulation. Further, our data show that class I COs are not dependent on CDK2 since the residual MutLγ foci are found in the complete absence of CDK2 in males. To our knowledge, this is the first study to (a) describe a meiotic phenotype that differentially affects the localization and dynamics of licensing and pro-CO/designation factors that are critical for appropriate CO patterning and (b) reveal that defects in CO licensing by impaired MutSγ function can alter the dynamic accumulation and exchange of DSB repair intermediate-associated CDKs, namely the transition from CDK4 predominance to CDK2.

Studies in budding yeast identified MSH4 and MSH5 as early members of the ZMM protein family, where they function primarily to promote crossover formation through the major class I pathway ^38–40^. However, in this organism, the number of MutSγ foci detected during prophase I is around half of the final number of total crossovers (approximately 45-50 *Msh4* foci compared to 90 COs per cell), but this frequency of *Msh4* events is only half of the total number of MutLγ-dependent COs ^15,80–82^, suggesting that the ZMM role of MutSγ does not account fully for the final repertoire of class I COs. By contrast, in *C. elegans*, MSH-5 foci are clearly reduced from zygonema to pachynema; however nematodes lack MutLγ, with MutSγ being the final marker of all six CO events ^78,83^. In mice, the earlier appearance of MutSγ foci relative to MutLγ foci, together with higher abundance of MutSγ foci and the more severe and earlier phenotypes observed in homozygous *Msh4* or *Msh5* mutants ^35,36,42^, led us to propose that mammalian MutSγ functions beyond the ZMM-driven class I CO designation to coordinate DNA damage repair across multiple pathways. Consistent with this model, a point mutant mouse allele of *Msh5* that disrupts the ATPase domain of the protein results in the complete loss of chiasmata at diakinesis, rather than a selective loss of those arising from the canonical class I pathway ^42^. With the addition of the current data showing for the first time that pro-CO protein dynamics can be separated from the dynamic exchange of CDK4 with CDK2 as a marker of the transition from licensing to designation, we add important new information regarding species-specific nuances in MutSγ function.

In seeking an explanation for the role of the mammalian-specific 38-amino acid C-terminal domain of MSH5, we have examined previous reports describing the regulation of MutSγ function through post-translational modification (PTM) across eukaryotes. In *S. cerevisiae*, for example, the N-terminus contains a degron that renders the protein intrinsically unstable and targets it for proteasomal degradation. Stabilization of MSH4 has been shown to depend on a phosphorylation event within this region ^80^. Coupled with this, two MSH4 sumoylation events are required for MutSγ formation and meiotic crossover formation in budding yeast ^84^. Consistent with the post-translational regulation of MutSγ in budding yeast, recent work in *C. elegans* has demonstrated that CDK-2 targets MutSγ by phosphorylating the disordered C-terminal tail of MSH-5, thereby stabilizing crossover-specific recombination intermediates ^70^. Interestingly, the C-terminal deletion of MSH5 results in abnormal retention of RNF212B and CDK4 through until pachynema in *Msh5*^ΔC*/*Δ^*^C^* male and female meiocytes, two potential modifiers of MSH5 via sumoylation and phosphorylation, respectively, while *in silico* analysis of the C-terminal domain reveals at least two predicted sites of PTMs, including phosphorylation and sumoylation domains. These data suggest a reciprocal relationship between MutSγ and PTM-invoking CO licensing regulators.

When comparing the phenotypic consequence of C-terminal deletion in *Msh5*^ΔC*/*Δ^*^C^* males and females, we observe a greater number of MutLγ foci in spermatocytes compared to oocytes, suggesting a degree of sexual dimorphism in CO designation efficiency. Recently, Horan *et al* studied the efficiency of C57Bl6/J male and female mice for establishing CO intermediates (licensing), conversion of DSBs to mature class I COs, and CO designation efficiency by analyzing the ratios of MSH4:RAD51 foci, MLH1:RAD51 foci and MLH1:MSH4 foci respectively. Although both males and females have the same licensing efficiency (MSH4:RAD51 ratios), males showed consistently higher efficiency to convert mature DSBs and MSH4 foci into mature class I COs ^71^. Along these lines, we observe that when the C-terminal domain of MSH5 is lost and the ability to ensure stabilization of MutSγ in the SC by pachynema is compromised, males are more efficient in converting these fewer foci into MutLγ foci than females, supporting these previous observations.

Studies of female meiotic mutants show that the timing of meiotic defects during prophase I strongly influences reproductive outcomes. Female mice lacking *Msh4*, *Msh5, Rnf212b*, and *Dmc1* exhibit meiotic disruption from zygonema, leading to a gradual depletion of ovarian follicles (and their contained oocytes) from shortly after birth and continuing for the first three weeks of life ^35,36,41,54,85,86^. By contrast, homozygous mutations in later CO designation genes (*Mlh1*, *Mlh3*, and *Cntd1*) allow oocytes to complete early prophase I and enter dictyate arrest, with defects becoming apparent only after ovulation and fertilization when meiosis resumes ^87^. Our analysis of *Msh5^ΔC/ΔC^* females reveals complete oocyte loss by PND1, indicating that elimination occurs before dictyate arrest and primordial follicle establishment. Notably, this loss occurs slightly earlier than in complete null mutants of MutSγ components, despite more advanced meiotic progression. Most likely, these germ cells are being eliminated from the ovary of *Msh5*^ΔC*/*Δ^*^C^* females through activation of the ATR-CHK2 signaling pathway, triggering p53/TAp63-dependent apoptosis before dictyate arrest ^88,89^. Our data suggest that defective meiocytes in both sexes are to be eliminated around the diplotene-like stage or slightly earlier, as no diakinesis-stage spermatocytes were detected in *Msh5*^ΔC*/*Δ^*^C^* males when performing chiasmata preparations.

Taken together, we propose that full-length MSH5 is essential for the full-spectrum function of MutSγ in mammals, both in promoting class I CO events and in supporting the repair of all DSB intermediates that persist after zygonema through multiple repair pathways. The complete loss of CDK2 association, together with the persistence of CDK4 into diplonema, suggests a mechanism in which MutSγ directly or indirectly enables the stepwise recruitment of distinct cyclin-dependent kinases, each likely targeting different substrates for phosphorylation. Additionally, we find that the association of CROs usually associated with licensing-to-designation steps are disconnected from these events. These dynamic associations of CDKs and CROs with nascent COs through prophase I is likely to regulate critical stepwise patterning of crossovers across the mammalian genome, driven in part by the diverse function of MutSγ through meiotic prophase I in mammals.

## RESOURCE AVAILABILITY

### Lead Contact

Paula E. Cohen

### Materials Availability

antibodies and mice are available upon request (email Lead Contact)

### Data and Code Availability

Not applicable

## ACKNOWLEDGEMENTS

We are indebted to the Cornell Center for Animal Resources and Education (CARE) for their assistance with mouse husbandry, to Eileen Shu for laboratory management, and to the members of the Cohen Lab for their contributions to this manuscript. We also thank Alberto Pendas for the RNF212B antibody and Mary Ann Handel and Ewelina Bolcun-Filas for the H1T antibody. The work described in this manuscript was supported by financial support from the Eunice Kennedy Shriver National Institute of Child Health and Development to PEC (R01HD041012 and R01HD097987) and by the National Cancer Institute to WE (R01CA248536).

## AUTHOR CONTRIBUTIONS

### Experimental Design

Paula E. Cohen, Winfried Edelmann, Ky’ara D. Carr, Maria de las Mercedes Carro.

## Experiments

Ky’ara D. Carr, Maame S. Adjei-Boadu, Eliza O’Donnell, Tegan S. Horan, Anna J. Wood, Yongwei Zhang, Maria de las Mercedes Carro.

### Data Analysis and Interpretation

Ky’ara D. Carr, Maame S. Adjei-Boadu, Eliza O’Donnell, Maria de las Mercedes Carro, Paula E. Cohen.

### Funding acquisition

Paula E. Cohen, Winfried Edelmann.

### Manuscript Preparation

Ky’ara D. Carr, Maria de las Mercedes Carro, Paula E. Cohen.

## MATERIALS AND METHODS

### Care and use of experimental animals

All mice used in this study were maintained on a C57BL/6J background acquired from The Jackson Laboratory. Mice were housed under strictly defined conditions of constant temperature, 12-hour light: dark cycles, provided with food and water *ad libitum*. All adult mice in this study were euthanized between 10 and 11 weeks of age using carbon dioxide, followed by cervical dislocation. All fetal mice in this study were euthanized at 18.5 days post coitum (dpc) by decapitation. Homozygous *Msh5* ^ΔC/ΔC^ and *Msh5* ^-/-^ animals were obtained through intercrosses of heterozygous animals. All mouse lines generated in these studies were backcrossed more than 10 times onto a C57BL/6J genetic background. Natural mating assays were conducted by housing males and females together until each breeding pair produced at least three litters. The litter size and offspring sex ratio were recorded and analyzed. Animal handling and procedures were performed following approval by the Cornell Institutional Animal Care and Use Committee, under protocol 2004-0063.

### Generation of mice and genotyping

#### A. Generation of *Msh5*^ΔC^ mice

Deletion of the MSH5 C-terminal domain was generated by inserting a premature stop codon at residue 795, positioned in exon 23 of the *Msh5* gene ^42^. The targeting vector pMsh5ex23 (50 μg) was electroporated into WW6 embryonic stem (ES) cells, and hygromycin–resistant colonies were isolated. Positive ES cell colonies were identified by PCR. The wild-type allele was detected as 600 bp, whereas the mutant allele was detected at 700 bp. Sanger sequencing of the PCR products was performed to verify the mutant allele. In silico analyses indicated that the deleted region contains at least two putative post-translational modification sites.

#### B. Generation of *Msh5*^null^ mice

Deletion of the *Msh5* gene was generated using the CRISPR-Cas9 system, in which a point mutation was introduced within the *Msh*5 genomic locus. The wild-type allele was detected as 387 bp, similar to the size of the null allele; thus, identification of the mutant allele was done by Sanger sequencing of the PCR products.

#### C. Genotyping of *Msh5*^ΔC^ and *Msh5*^null^ mice

Genomic DNA was extracted from all mice used in this study from ear snips by a 20-minute incubation at 95°C in 100 μL of alkaline solution (0.05M NaOH), followed by pH neutralization with 10 μL of 1M Tris-HCl, pH 7.2, and finally mixed by pulse vortexing. All genotyping was performed using 10μM primers and OneTaq 2X Master Mix (NEB M0482) according to the manufacturer’s instructions.

##### PCR conditions for *Msh5* ^ΔC^ line

PCR amplification of both wild-type and mutant alleles was performed under the following conditions: an initial denaturation at 94°C for 3 min; 40 cycles of 95°C for 20 s, 60°C for 30 s, 72°C for 1 min; followed by a final extension at 72°C for 10 min, and a hold at 4°C. Genotyping was performed using the following primers: Msh5delC-F (5’-TGAGGATGGGGAAGACCTTG -3’) and Msh5delC-R (5’-AGGGAGAACACAGGAGCTCT-3’). The wild-type and mutant alleles yielded PCR products of 600 and 700 bp, respectively.

##### PCR conditions for *Msh5* ^null^ line

PCR amplification of both wild-type and mutant alleles was performed under the following conditions: an initial denaturation a**t** 95°C for 2.5 min; 32 cycles of 95°C for 20 secs, 56°C for 30 secs, 72°C for 1 min; followed by a final extension at 72°C for 6 min, and a hold at 4°C. Genotyping was performed using the following primers: Msh5KO-F (5’-CACGTTGACTAAACTCCCCA -3’) and Msh5KO-R, (5’-TCACATACAGTAGGTACCTA -3’). Subsequently, the PCR product is purified using the QIAquick PCR Purification Kit and followed by Sanger sequencing.

#### Epididymal Sperm Counts

Bilateral cauda epididymides were collected from mice and minced with scalpel blades in 1-mL of pre-warmed Dulbecco’s Modified Eagle Medium (DMEM) supplemented with 4% bovine serum albumin (BSA), and incubated at 37°C with 5% CO_2_ for 20 min to allow spermatozoa to swim out of each epididymis. An aliquot (20 μL) of the sperm suspension was fixed in 480 μL of 10% formalin, and sperm counts were obtained using a hemacytometer.

#### Histological analysis of mouse testis and ovaries

Adult testes were fixed in Bouin’s solution or 10% formalin for 6 hours at room temperature, and then washed in 70% ethanol. Fixed testes were embedded in paraffin and sectioned at 5μm. Ovaries from mice at postnatal days 1, 5, and 10-week-old adults were fixed in either Bouin’s or formalin for 4 hours at room temperature and then washed in 70% ethanol. All ovarian tissues were sectioned and paraffin-embedded in 5-micron thick sections by the Cornell Histopathology core. Tissues were deparaffinized as follows: three 10-minute washes in Safeclear, followed by three 3-minute washes in 95% ethanol, subsequently followed by one 5-minute wash in 80% ethanol and 70% ethanol, with a final wash in 1X PBS for 10 minutes. Staining comprised of one 30-second staining in Hematoxylin followed by two washes in miliQ-H_2_O, followed by two dunks in Bluing reagent and subsequent washes in miliQ-H_2_O. Slides were stained with Eosin for 2 minutes followed by four dunks in miliQ-H_2_O. Slides were then processed through 1 dunk of 50% ethanol, 70% ethanol and 80% ethanol, and three 3-minute washes in 95% ethanol, finished with three 3-minute Safeclear washes. Slides were dried, mounted with Permount, and subsequently imaged using an Aperio CS2 Digital Pathology Slide Scanscope. The total number of follicles was determined by multiplying the raw counts from every fifth section by 5 to correct for the sections not counted. Follicles were staged as explained previously ^51,90^, and follicles with oocytes without a visible nucleus were excluded from quantification to avoid double-counting.

#### Spermatocyte chromosome preparations and immunofluorescence staining

Meiotic spreads were processed from 10-week-old adult testes according to the previously described method ^50,52,91,92^. Briefly, testes were detunicated in phosphate-buffered saline (PBS), and tubules were incubated in hypotonic extraction buffer (50 mM sucrose; 17 mM Citrate; 30 mM Tris; 5 mM EDTA; 0.5 mM DTT; 0.1 mM PMSF; pH 8.2-8.4) for 2 hours on ice. Seminiferous tubules from each testis were segmented and minced in 100 mM sucrose at room temperature. Cell suspensions were dispersed onto slides coated in 1% paraformaldehyde (pH 9.2) containing 0.15% Triton X-100. Slides were incubated overnight in a humid chamber at room temperature and then air-dried for 30 minutes. Slides were washed with 0.4% Kodak Photo-Flo 200 solution, and then air dried completely, and then stored at -80°C until use or immediately stained. For staining, slides are washed for 10 minutes in PBS with 0.4% Photo-Flo, followed by PBS with 0.1% Triton X-100 for 10 minutes, and blocked for 30 minutes in 10% antibody dilution buffer (PBS; 3% bovine serum albumin; 10% goat serum; Triton X-100; filter sterilized). Primary antibodies were diluted in antibody dilution buffer at varying concentrations and applied to slides (MSH4 [rabbit] 1:100, RAD51 1:500, SYCP3 [mouse] 1:1000, SYCP3 [rabbit] 1:1,000, γH2AX [mouse] 1:10,000, HORMAD1 [rabbit] 1:1,000, SYCP1 [rabbit] 1:1,000, MLH1 [mouse] 1:50, MLH3 [guinea pig] 1:500, RNF212B [rabbit] 1:750, HEI10 [rabbit] 1:100, CNTD1 [rabbit] 1:100, CDK2 [mouse] 1:50, CDK4 [rabbit] 1:100, H1T [guinea pig] 1:500), followed by an overnight incubation in a humid chamber at room temperature. Slides are washed for 10 minutes in PBS with 0.4% Photo-Flo, followed by 10 minutes in PBS with 0.1% Triton X-100, and finally blocked for 10 minutes in 10% antibody dilution buffer. Secondary antibodies were diluted in antibody dilution buffer and then subsequently applied to slides and incubated in a humid chamber for 1 hour at 37°C. Slides were washed three times for 5 minutes in PBS with 0.4% Photo-Flo, followed by 5 minutes in Mili-Q water with 0.4% Photo-Flo, and then air-dried for 45 minutes. Glass coverslips were mounted with 75 μL of Prolong Antifade with DAPI, applied to the slides, and then sealed with clear nail polish.

#### Diakenesis spread preparation for the observation of chiasmata

Spermatocyte diakinesis spreads were prepared from 10-week-old adult testes according to the previously described method ^52,91–93^. Briefly, testes were detunicated and transferred onto a depression slide with 2.2% sodium citrate (pre-warmed at 37°C). Tubules were minced with a scalpel blade, followed by further dissociation of the tubules with a Pasteur pipette. Tubule fragments were left to settle for 15 minutes at room temperature, and then the cell suspension was collected for centrifugation at 200xg for 5 minutes. The pellets formed were resuspended in 4 mL of pre-warmed 1.1% sodium citrate and incubated at room temperature for 15 minutes. The cell suspension was centrifuged at 200xg for 5 minutes, and the supernatant was discarded. The cell pellet was resuspended vigorously in fixative (3 parts of methanol: 1 part of glacial acetic acid), followed by a 5-minute incubation at room temperature. The fixed suspension was centrifuged at 200xg for 5 minutes, and then the fixative was discarded. Before slide preparation, the fixed cells underwent two additional rounds of centrifugation and room temperature incubations. Slides were immersed in fixative for 15 minutes before use. Small drops of cell suspension were applied to slides and air dried. Slides were stained with 4% Giemsa (Fisher) for 6 minutes, followed by three washes for 3 minutes in double-distilled water, and finally mounted with permount.

#### Oocyte chromosome preparations and immunofluorescence staining

Oocytes were processed from mouse fetal ovaries isolated at 18.5 days post coitum (dpc), with 0.5 dpc defined by the detection of a vaginal plug, and oocytes were subsequently processed as previously described ^49,51,94^. Fetal ovaries were transferred into PBS, followed by incubation in hypotonic extraction buffer for 20 - 30 minutes at room temperature. Fetal ovaries were minced in 100 mM sucrose at room temperature, and then the cell suspension was applied to slides coated in 1% paraformaldehyde (pH 9.2) containing 0.15% Triton X-100. Slides were incubated in a humid chamber at room temperature for 2 hours and then air-dried at room temperature. Primary antibodies were diluted in antibody dilution buffer at varying concentrations (SYCP3 [mouse] 1:500). Other slides were stained and mounted as described for spermatocyte chromosome spreads.

### Image Acquisition

All immunostained slides were kept at 4°C until imaging on an epifluorescence microscope (Axiophot Imager Z1) equipped with Zeiss Zen Blue version 3.0 software (Carl Zeiss AG, Oberkochen, Germany). Images were taken under 63X magnification and processed using the Fiji software. For each applied antibody, standard conditions were consistent in exposure times. SYCP3, MLH3, and H1T were used to stage prophase I cells. The extent of X-Y chromosome synapsis was used to assign spermatocytes to early, mid, or late pachynema, as previously described ^95^. Pachynema-like meiocytes were defined by at least 10 fully synapsed SCs. Cells were designated as “diplotene-like” when they displayed bulging ends, desynapsis, and/or reduced length of the pseudoautosomal region (PAR). Slide stained with Giesma was imaged using brightfield under Zeiss Zen Blue version 3.0 software (Carl Zeiss AG, Oberkochen, Germany). Images were taken under 40X magnification and then processed as described for immunostained slides.

### Statistical methods and analysis

All data were analyzed and interpreted using GraphPad Prism version 10.4.1. Testis weights and cauda epididymal sperm counts were analyzed using an unpaired nonparametric Mann-Whitney test. Mann-Whitney tests were also used to compare MSH4, RAD51, MLH1, and MLH3 foci per nucleus in wild-type and mutant male and female mice. At least 3 animals were analyzed per group. All p-values that are less than 0.05 in this study were considered significantly different.

## SUPPLEMENTAL INFORMATION

### Supplemental Figure Legends

**Figure S1. Schematic representation of the *Msh5*^ΔC^ mouse design strategy.** (A) Embryonic stem cells carrying the targeting vector were utilized to generate the *Msh5*^ΔC^ mutant mouse line ^42^. The floxed allele in the male breeder showed the PGK hygromycin cassette, and the female mice expressed a Cre recombinase controlled by the zona pellucida (Zp3) gene promoter. (B) PCR amplification from genomic DNA generated products of 600 bp and 700 bp from *Msh5^+/+^* and *Msh5^ΔC/ΔC^*, respectively. A representative genomic sequence trace from *Msh5^ΔC/ΔC^* is shown, demonstrating deletion of the C-terminal domain encoding 38 amino acids (aa). The *Msh5^ΔC^* mouse harbors the following in-frame mutation in exon 23 (indicated by white arrow). (C) Multiple sequence alignment of the MSH5 C-terminal domain was performed using Clustal Omega across diverse species, including budding yeast, plants, nematodes, zebrafish, chicken, koala, mouse, rat, elephant, human, and bovine. The magenta box highlights a conserved region shared among all mammalian species examined.

**Figure S2. Number of pups and litter size** (A) Number of offspring per litter produced by natural mating. (B) Log_2_ (female/male pup ratio) after natural mating. Dots represent the number of pups per liter, and bars represent the average of pups ± SD, n ≥ 3. Statistical comparisons were performed using either (A) one-way ANOVA or (B) the Mann-Whitney test; p-values are either shown as p < 0.0001 (****) or ns = not significant.

**Figure S3. Autosomal flares of γH2AX signal are detected in pachytene-staged *Msh5*** ^Δ**C/**Δ**C**^ **spermatocytes.** (A-H) Representative images of spermatocyte prophase I spreads from *Msh5^+/+^* and *Msh5^ΔC/ΔC^* male mice immunostained against γH2AX (green, proxy for DNA damage) and SYCP3 (magenta) in (A-D) *Msh5*^+/+^ and (E-H) *Msh5*^ΔC/ΔC^ adult males. White scale bars are equal to 10μm.

**Figure S4. *Msh5***^Δ**C/**Δ**C**^ **spermatocytes exhibit synaptic defects and non-homologous interactions.** (A-F) Representative images of pachytene and pachytene-like spermatocyte prophase I spreads from *Msh5^+/+^*and *Msh5^ΔC/ΔC^* male mice immunostained against SYCP1 (green, transverse filament of the SC) and SYCP3 (magenta) in (A-D) *Msh5*^+/+^ and (E-H) *Msh5*^ΔC/ΔC^ adult males. (G-H) Representative images of pachytene and pachytene-like spermatocyte prophase I spreads from *Msh5^+/+^*and *Msh5^ΔC/ΔC^* male mice immunostained against HORMAD1 (green) and SYCP3 (magenta), from *Msh5*^+/+^ and *Msh5*^ΔC/ΔC^ adult males. White scale bars are equal to 10μm.

**Figure S5.** Unrepaired DSBs and Autosomal flares of γH2AX signal and synaptic defects are detected in pachytene-staged *Msh5*^Δ**C/**Δ**C**^ **oocytes.** (A-F) Representative images of oocyte prophase I spreads from *Msh5^+/+^* and *Msh5^ΔC/ΔC^* female mice immunostained against γH2AX (green, proxy for DNA damage) and SYCP3 (magenta). White scale bars are equal to 10μm. (G-L) Representative images of oocyte prophase I spreads from *Msh5^+/+^*and *Msh5 ^ΔC/ΔC^* 18.5 dpc female embryos immunostained against RAD51 (green) as a proxy for DSB repair and SYCP3 (magenta). White scale bars are equal to 10μm. (M) Quantification of total RAD51 foci per pachytene and pachytene-like nucleus. Each point represents the average of foci per nucleus, and bars represent the average foci number ± SD, n = 16 cells from 3 mice. Statistical comparisons were performed using the Mann-Whitney test; p-values are either shown as p < 0.0001 (****) or ns = not significant. (N-S) Representative images of pachytene and pachytene-like oocyte prophase I spreads from *Msh5^+/+^* and *Msh5^ΔC/ΔC^* 18.5 female embryos immunostained against SYCP1 (green, transverse filament of the SC) and SYCP3 (magenta). (T-U) Representative images of pachytene and pachytene-like oocyte prophase I spreads from *Msh5^+/+^* and *Msh5^ΔC/ΔC^* 18.5 female embryos immunostained against HORMAD1 (green) and SYCP3 (magenta). White scale bars are equal to 10μm. Arrows indicate inter-homologue interactions.

**Figure S6. *Msh5^ΔC/ΔC^* oocytes have impaired localization of pro-crossover factors and cyclin-dependent kinases.** Representative images of pachytene and pachytene-like oocyte prophase I spreads from *Msh5^+/+^*and *Msh5^ΔC/ΔC^* female embryos immunostained against RNF212B (A-B, G-H), HEI10 (C-E, I-K), CNTD1 (F, L), CDK4 (M-P, R-U), and CDK2 (Q, V) in green and SYCP3 in magenta, cells from 3 mice. White scale bars are equal to 10 μm. Arrows indicate telomeric (red) and CO-associated (white) CDK2.

## REFERENCES

1. Wang, S., and Zhang, L. (2026). Meiotic crossover control: Interplay between the recombination process and chromosome organization. Curr. Top. Dev. Biol. 168, 281–315. 10.1016/bs.ctdb.2026.01.003.

2. Yang, F., and Wang, P.J. (2009). The Mammalian synaptonemal complex: a scaffold and beyond. Genome Dyn. 5, 69–80. 10.1159/000166620.

3. Jiang, H., Fan, S., and Shi, Q. (2026). Synaptonemal complex: The structural basis for meiosis I. Curr. Top. Dev. Biol. 168, 245–279. 10.1016/bs.ctdb.2026.01.014.

4. Cesar, B.I., and Kim, Y. (2026). Structure and function of the synaptonemal complex. J. Cell Biol. 225. 10.1083/jcb.202511222.

5. Zheng, Z., Zheng, L., Arter, M., Liu, K., Yamada, S., Ontoso, D., Kim, S., and Keeney, S. (2024). Reconstitution of SPO11-dependent double-strand break formation. BioRxiv. 10.1101/2024.11.20.624382.

6. Keeney, S., Baudat, F., Angeles, M., Zhou, Z.H., Copeland, N.G., Jenkins, N.A., Manova, K., and Jasin, M. (1999). A mouse homolog of the Saccharomyces cerevisiae meiotic recombination DNA transesterase Spo11p. Genomics 61, 170–182. 10.1006/geno.1999.5956.

7. Baudat, F., Manova, K., Yuen, J.P., Jasin, M., and Keeney, S. (2000). Chromosome synapsis defects and sexually dimorphic meiotic progression in mice lacking *Spo11*. Mol. Cell 6, 989–998. 10.1016/S1097-2765(00)00098-8.

8. Keeney, S., Giroux, C.N., and Kleckner, N. (1997). Meiosis-specific DNA double-strand breaks are catalyzed by Spo11, a member of a widely conserved protein family. Cell 88, 375–384. 10.1016/s0092-8674(00)81876-0.

9. Tang, X., and Tong, M.-H. (2026). Mechanism and regulation of meiotic double-strand break formation in mammals. Trends Biochem. Sci. 51, 354–366. 10.1016/j.tibs.2026.01.006.

10. Hassold, T., Hall, H., and Hunt, P. (2007). The origin of human aneuploidy: where we have been, where we are going. Hum. Mol. Genet. 16 *Spec No. 2*, R203-8. 10.1093/hmg/ddm243.

11. Lenzi, M.L., Smith, J., Snowden, T., Kim, M., Fishel, R., Poulos, B.K., and Cohen, P.E. (2005). Extreme heterogeneity in the molecular events leading to the establishment of chiasmata during meiosis i in human oocytes. Am. J. Hum. Genet. 76, 112–127. 10.1086/427268.

12. Fishel, R. (2015). Mismatch repair. J. Biol. Chem. 290, 26395–26403. 10.1074/jbc.R115.660142.

13. Hunter, N., and Borts, R.H. (1997). Mlh1 is unique among mismatch repair proteins in its ability to promote crossing-over during meiosis. Genes Dev. 11, 1573–1582. 10.1101/gad.11.12.1573.

14. Ross-Macdonald, P., and Roeder, G.S. (1994). Mutation of a meiosis-specific MutS homolog decreases crossing over but not mismatch correction. Cell 79, 1069–1080. 10.1016/0092-8674(94)90037-x.

15. Novak, J.E., Ross-Macdonald, P.B., and Roeder, G.S. (2001). The budding yeast Msh4 protein functions in chromosome synapsis and the regulation of crossover distribution. Genetics 158, 1013–1025.

16. Wang, T.F., Kleckner, N., and Hunter, N. (1999). Functional specificity of MutL homologs in yeast: evidence for three Mlh1-based heterocomplexes with distinct roles during meiosis in recombination and mismatch correction. Proc Natl Acad Sci USA 96, 13914–13919. 10.1073/pnas.96.24.13914.

17. Pochart, P., Woltering, D., and Hollingsworth, N.M. (1997). Conserved properties between functionally distinct MutS homologs in yeast. J. Biol. Chem. 272, 30345–30349. 10.1074/jbc.272.48.30345.

18. Colas, I., Macaulay, M., Higgins, J.D., Phillips, D., Barakate, A., Posch, M., Armstrong, S.J., Franklin, F.C.H., Halpin, C., Waugh, R., et al. (2016). A spontaneous mutation in MutL-Homolog 3 (HvMLH3) affects synapsis and crossover resolution in the barley desynaptic mutant des10. New Phytol. 212, 693–707. 10.1111/nph.14061.

19. Mao, B., Zheng, W., Huang, Z., Peng, Y., Shao, Y., Liu, C., Tang, L., Hu, Y., Li, Y., Hu, L., et al. (2021). Rice MutLγ, the MLH1-MLH3 heterodimer, participates in the formation of type I crossovers and regulation of embryo sac fertility. Plant Biotechnol. J. 19, 1443–1455. 10.1111/pbi.13563.

20. Capilla-Perez, L., Solier, V., Portemer, V., Chambon, A., Hurel, A., Guillebaux, A., Vezon, D., Cromer, L., Grelon, M., and Mercier, R. (2018). The HEM Lines: A New Library of Homozygous Arabidopsis thaliana EMS Mutants and its Potential to Detect Meiotic Phenotypes. Front. Plant Sci. 9, 1339. 10.3389/fpls.2018.01339.

21. Lipkin, S.M., Moens, P.B., Wang, V., Lenzi, M., Shanmugarajah, D., Gilgeous, A., Thomas, J., Cheng, J., Touchman, J.W., Green, E.D., et al. (2002). Meiotic arrest and aneuploidy in MLH3-deficient mice. Nat. Genet. 31, 385–390. 10.1038/ng931.

22. Hunter, N. (2015). Meiotic recombination: the essence of heredity. Cold Spring Harb. Perspect. Biol. 7. 10.1101/cshperspect.a016618.

23. Gray, S., and Cohen, P.E. (2016). Control of meiotic crossovers: from double-strand break formation to designation. Annu. Rev. Genet. 50, 175–210. 10.1146/annurev-genet-120215-035111.

24. Sidhu, G.K., Fang, C., Olson, M.A., Falque, M., Martin, O.C., and Pawlowski, W.P. (2015). Recombination patterns in maize reveal limits to crossover homeostasis. Proc Natl Acad Sci USA 112, 15982–15987. 10.1073/pnas.1514265112.

25. Cole, F., Kauppi, L., Lange, J., Roig, I., Wang, R., Keeney, S., and Jasin, M. (2012). Homeostatic control of recombination is implemented progressively in mouse meiosis. Nat. Cell Biol. 14, 424–430. 10.1038/ncb2451.

26. Dai, J., Sanchez, A., Adam, C., Ranjha, L., Reginato, G., Chervy, P., Tellier-Lebegue, C., Andreani, J., Guérois, R., Ropars, V., et al. (2021). Molecular basis of the dual role of the Mlh1-Mlh3 endonuclease in MMR and in meiotic crossover formation. Proc Natl Acad Sci USA 118. 10.1073/pnas.2022704118.

27. Toledo, M., Sun, X., Brieño-Enríquez, M.A., Raghavan, V., Gray, S., Pea, J., Milano, C.R., Venkatesh, A., Patel, L., Borst, P.L., et al. (2019). A mutation in the endonuclease domain of mouse MLH3 reveals novel roles for MutLγ during crossover formation in meiotic prophase I. PLoS Genet. 15, e1008177. 10.1371/journal.pgen.1008177.

28. Sonntag Brown, M., Lim, E., Chen, C., Nishant, K.T., and Alani, E. (2013). Genetic analysis of mlh3 mutations reveals interactions between crossover promoting factors during meiosis in baker’s yeast. G3 (Bethesda) 3, 9–22. 10.1534/g3.112.004622.

29. Flores-Rozas, H., and Kolodner, R.D. (1998). The Saccharomyces cerevisiae MLH3 gene functions in MSH3-dependent suppression of frameshift mutations. Proc Natl Acad Sci USA 95, 12404–12409. 10.1073/pnas.95.21.12404.

30. Cannavo, E., Sanchez, A., Anand, R., Ranjha, L., Hugener, J., Adam, C., Acharya, A., Weyland, N., Aran-Guiu, X., Charbonnier, J.-B., et al. (2020). Regulation of the MLH1-MLH3 endonuclease in meiosis. Nature 586, 618–622. 10.1038/s41586-020-2592-2.

31. Paquis-Flucklinger, V., Santucci-Darmanin, S., Paul, R., Saunières, A., Turc-Carel, C., and Desnuelle, C. (1997). Cloning and expression analysis of a meiosis-specific MutS homolog: the human MSH4 gene. Genomics 44, 188–194. 10.1006/geno.1997.4857.

32. Frasca, M., Paniker, L., Kang, R., Chakraborty, P., Pandey, A., LoPresti, J., and Cole, F. (2025). MutSgamma promotes meiotic recombination and homolog pairing in mouse spermatocytes. Genetics. 10.1093/genetics/iyaf099.

33. Higgins, J.D., Armstrong, S.J., Franklin, F.C.H., and Jones, G.H. (2004). The Arabidopsis MutS homolog AtMSH4 functions at an early step in recombination: evidence for two classes of recombination in Arabidopsis. Genes Dev. 18, 2557–2570. 10.1101/gad.317504.

34. Higgins, J.D., Vignard, J., Mercier, R., Pugh, A.G., Franklin, F.C.H., and Jones, G.H. (2008). AtMSH5 partners AtMSH4 in the class I meiotic crossover pathway in Arabidopsis thaliana, but is not required for synapsis. Plant J. 55, 28–39. 10.1111/j.1365-313X.2008.03470.x.

35. Kneitz, B., Cohen, P.E., Avdievich, E., Zhu, L., Kane, M.F., Hou, H., Kolodner, R.D., Kucherlapati, R., Pollard, J.W., and Edelmann, W. (2000). MutS homolog 4 localization to meiotic chromosomes is required for chromosome pairing during meiosis in male and female mice. Genes Dev. 14, 1085–1097. 10.1101/gad.14.9.1085.

36. Edelmann, W., Cohen, P.E., Kneitz, B., Winand, N., Lia, M., Heyer, J., Kolodner, R., Pollard, J.W., and Kucherlapati, R. (1999). Mammalian MutS homologue 5 is required for chromosome pairing in meiosis. Nat. Genet. 21, 123–127. 10.1038/5075.

37. Kolas, N.K., Svetlanov, A., Lenzi, M.L., Macaluso, F.P., Lipkin, S.M., Liskay, R.M., Greally, J., Edelmann, W., and Cohen, P.E. (2005). Localization of MMR proteins on meiotic chromosomes in mice indicates distinct functions during prophase I. J. Cell Biol. 171, 447–458. 10.1083/jcb.200506170.

38. Shinohara, M., Oh, S.D., Hunter, N., and Shinohara, A. (2008). Crossover assurance and crossover interference are distinctly regulated by the ZMM proteins during yeast meiosis. Nat. Genet. 40, 299–309. 10.1038/ng.83.

39. Lynn, A., Soucek, R., and Börner, G.V. (2007). ZMM proteins during meiosis: crossover artists at work. Chromosome Res. 15, 591–605. 10.1007/s10577-007-1150-1.

40. Börner, G.V., Kleckner, N., and Hunter, N. (2004). Crossover/noncrossover differentiation, synaptonemal complex formation, and regulatory surveillance at the leptotene/zygotene transition of meiosis. Cell 117, 29–45. 10.1016/s0092-8674(04)00292-2.

41. de Vries, S.S., Baart, E.B., Dekker, M., Siezen, A., de Rooij, D.G., de Boer, P., and te Riele, H. (1999). Mouse MutS-like protein Msh5 is required for proper chromosome synapsis in male and female meiosis. Genes Dev. 13, 523–531. 10.1101/gad.13.5.523.

42. Milano, C.R., Holloway, J.K., Zhang, Y., Jin, B., Smith, C., Bergman, A., Edelmann, W., and Cohen, P.E. (2019). Mutation of the ATPase Domain of MutS Homolog-5 (MSH5) Reveals a Requirement for a Functional MutSγ Complex for All Crossovers in Mammalian Meiosis. G3 (Bethesda) 9, 1839–1850. 10.1534/g3.119.400074.

43. Anderson, L.K., Reeves, A., Webb, L.M., and Ashley, T. (1999). Distribution of crossing over on mouse synaptonemal complexes using immunofluorescent localization of MLH1 protein. Genetics 151, 1569–1579.

44. Santucci-Darmanin, S., Walpita, D., Lespinasse, F., Desnuelle, C., Ashley, T., and Paquis-Flucklinger, V. (2000). MSH4 acts in conjunction with MLH1 during mammalian meiosis. FASEB J. 14, 1539–1547. 10.1096/fj.14.11.1539.

45. Baker, S.M., Plug, A.W., Prolla, T.A., Bronner, C.E., Harris, A.C., Yao, X., Christie, D.M., Monell, C., Arnheim, N., Bradley, A., et al. (1996). Involvement of mouse Mlh1 in DNA mismatch repair and meiotic crossing over. Nat. Genet. 13, 336–342. 10.1038/ng0796-336.

46. Holloway, J.K., Booth, J., Edelmann, W., McGowan, C.H., and Cohen, P.E. (2008). MUS81 generates a subset of MLH1-MLH3-independent crossovers in mammalian meiosis. PLoS Genet. 4, e1000186. 10.1371/journal.pgen.1000186.

47. Edelmann, W., Cohen, P.E., Kane, M., Lau, K., Morrow, B., Bennett, S., Umar, A., Kunkel, T., Cattoretti, G., Chaganti, R., et al. (1996). Meiotic pachytene arrest in MLH1-deficient mice. Cell 85, 1125–1134. 10.1016/s0092-8674(00)81312-4.

48. Svetlanov, A., Baudat, F., Cohen, P.E., and de Massy, B. (2008). Distinct functions of MLH3 at recombination hot spots in the mouse. Genetics 178, 1937–1945. 10.1534/genetics.107.084798.

49. Kan, R., Sun, X., Kolas, N.K., Avdievich, E., Kneitz, B., Edelmann, W., and Cohen, P.E. (2008). Comparative analysis of meiotic progression in female mice bearing mutations in genes of the DNA mismatch repair pathway. Biol. Reprod. 78, 462–471. 10.1095/biolreprod.107.065771.

50. Gray, S., Santiago, E.R., Chappie, J.S., and Cohen, P.E. (2020). Cyclin N-Terminal Domain-Containing-1 Coordinates Meiotic Crossover Formation with Cell-Cycle Progression in a Cyclin-Independent Manner. Cell Rep. 32, 107858. 10.1016/j.celrep.2020.107858.

51. Wood, A.J., Ahmed, R.M., Simon, L.E., Bradley, R.A., Gray, S., Wolff, I.D., and Cohen, P.E. (2025). CNTD1 is crucial for crossover formation in female meiosis and for establishing the ovarian reserve. J. Cell Biol. 224. 10.1083/jcb.202401021.

52. Holloway, J.K., Sun, X., Yokoo, R., Villeneuve, A.M., and Cohen, P.E. (2014). Mammalian CNTD1 is critical for meiotic crossover maturation and deselection of excess precrossover sites. J. Cell Biol. 205, 633–641. 10.1083/jcb.201401122.

53. Qiao, H., Prasada Rao, H.B.D., Yang, Y., Fong, J.H., Cloutier, J.M., Deacon, D.C., Nagel, K.E., Swartz, R.K., Strong, E., Holloway, J.K., et al. (2014). Antagonistic roles of ubiquitin ligase HEI10 and SUMO ligase RNF212 regulate meiotic recombination. Nat. Genet. 46, 194–199. 10.1038/ng.2858.

54. Condezo, Y.B., Sainz-Urruela, R., Gomez-H, L., Salas-Lloret, D., Felipe-Medina, N., Bradley, R., Wolff, I.D., Tanis, S., Barbero, J.L., Sánchez-Martín, M., et al. (2024). RNF212B E3 ligase is essential for crossover designation and maturation during male and female meiosis in the mouse. Proc Natl Acad Sci USA 121, e2320995121. 10.1073/pnas.2320995121.

55. Ito, M., Yun, Y., Kulkarni, D.S., Lee, S., Sandhu, S., Nuñez, B., Hu, L., Lee, K., Lim, N., Hirota, R.M., et al. (2025). Distinct and interdependent functions of three RING proteins regulate recombination during mammalian meiosis. Proc Natl Acad Sci USA 122, e2412961121. 10.1073/pnas.2412961121.

56. Ashley, T., Walpita, D., and de Rooij, D.G. (2001). Localization of two mammalian cyclin dependent kinases during mammalian meiosis. J. Cell Sci. 114, 685–693. 10.1242/jcs.114.4.685.

57. Hamer, G., Roepers-Gajadien, H.L., van Duyn-Goedhart, A., Gademan, I.S., Kal, H.B., van Buul, P.P.W., and de Rooij, D.G. (2003). DNA double-strand breaks and gamma-H2AX signaling in the testis. Biol. Reprod. 68, 628–634.

58. Prabhu, K.S., Kuttikrishnan, S., Ahmad, N., Habeeba, U., Mariyam, Z., Suleman, M., Bhat, A.A., and Uddin, S. (2024). H2AX: A key player in DNA damage response and a promising target for cancer therapy. Biomed. Pharmacother. 175, 116663. 10.1016/j.biopha.2024.116663.

59. Mahadevaiah, S.K., Turner, J.M., Baudat, F., Rogakou, E.P., de Boer, P., Blanco-Rodríguez, J., Jasin, M., Keeney, S., Bonner, W.M., and Burgoyne, P.S. (2001). Recombinational DNA double-strand breaks in mice precede synapsis. Nat. Genet. 27, 271–276. 10.1038/85830.

60. Ashley, T., Plug, A.W., Xu, J., Solari, A.J., Reddy, G., Golub, E.I., and Ward, D.C. (1995). Dynamic changes in Rad51 distribution on chromatin during meiosis in male and female vertebrates. Chromosoma 104, 19–28.

61. Plug, A.W., Xu, J., Reddy, G., Golub, E.I., and Ashley, T. (1996). Presynaptic association of Rad51 protein with selected sites in meiotic chromatin. Proc Natl Acad Sci USA 93, 5920–5924.

62. de Vries, F.A.T., de Boer, E., van den Bosch, M., Baarends, W.M., Ooms, M., Yuan, L., Liu, J.-G., van Zeeland, A.A., Heyting, C., and Pastink, A. (2005). Mouse *Sycp1* functions in synaptonemal complex assembly, meiotic recombination, and XY body formation. Genes Dev. 19, 1376–1389. 10.1101/gad.329705.

63. Wojtasz, L., Daniel, K., Roig, I., Bolcun-Filas, E., Xu, H., Boonsanay, V., Eckmann, C.R., Cooke, H.J., Jasin, M., Keeney, S., et al. (2009). Mouse HORMAD1 and HORMAD2, two conserved meiotic chromosomal proteins, are depleted from synapsed chromosome axes with the help of TRIP13 AAA-ATPase. PLoS Genet. 5, e1000702. 10.1371/journal.pgen.1000702.

64. Milburn, A.E., Kulkami, D.S., Espejo-Serrano, C., Pachon-Penalba, M., Williams, M.E., Nicol, J.P.O., Debilio, S., Gurusaran, M., Dunce, J.M., Adams, I.R., et al. (2026). Molecular architecture of meiotic pro-crossover factor HEI10 reveals coupling of higher-order assembly and ubiquitin chain formation. BioRxiv. 10.64898/2026.05.08.723602.

65. Reynolds, A., Qiao, H., Yang, Y., Chen, J.K., Jackson, N., Biswas, K., Holloway, J.K., Baudat, F., de Massy, B., Wang, J., et al. (2013). RNF212 is a dosage-sensitive regulator of crossing-over during mammalian meiosis. Nat. Genet. 45, 269–278. 10.1038/ng.2541.

66. Ward, J.O., Reinholdt, L.G., Motley, W.W., Niswander, L.M., Deacon, D.C., Griffin, L.B., Langlais, K.K., Backus, V.L., Schimenti, K.J., O’Brien, M.J., et al. (2007). Mutation in mouse *Hei10*, an e3 ubiquitin ligase, disrupts meiotic crossing over. PLoS Genet. 3, e139. 10.1371/journal.pgen.0030139.

67. Tsutsui, T., Hesabi, B., Moons, D.S., Pandolfi, P.P., Hansel, K.S., Koff, A., and Kiyokawa, H. (1999). Targeted Disruption of CDK4 Delays Cell Cycle Entry with Enhanced p27^Kip1^ Activity. Mol. Cell. Biol. 19, 7011–7019. 10.1128/MCB.19.10.7011.

68. Palmer, N., Talib, S.Z.A., Singh, P., Goh, C.M.F., Liu, K., Schimenti, J.C., and Kaldis, P. (2020). A novel function for CDK2 activity at meiotic crossover sites. PLoS Biol. 18, e3000903. 10.1371/journal.pbio.3000903.

69. Bradley, R.A., Wolff, I.D., Cohen, P.E., and Gray, S. (2023). Dynamic regulatory phosphorylation of mouse CDK2 occurs during meiotic prophase I. BioRxiv. 10.1101/2023.07.24.550435.

70. Haversat, J., Woglar, A., Klatt, K., Akerib, C.C., Roberts, V., Chen, S.-Y., Arur, S., Villeneuve, A.M., and Kim, Y. (2022). Robust designation of meiotic crossover sites by CDK-2 through phosphorylation of the MutSγ complex. Proc Natl Acad Sci USA 119, e2117865119. 10.1073/pnas.2117865119.

71. Horan, T.S., Wood, A., Tanis, S., Gabarrell, C.P., and Cohen, P.E. (2026). Molecular assessment of recombination processing across genetically diverse 3. Mol Biol Evol.

72. Peterson, A.L., and Payseur, B.A. (2021). Sex-specific variation in the genome-wide recombination rate. Genetics 217, 1–11. 10.1093/genetics/iyaa019.

73. Peterson, A.L., and Payseur, B.A. (2021). Higher Intercellular Variation in Genome-Wide Recombination Rate in Female Mice. Cytogenet. Genome Res. 161, 463–469. 10.1159/000516998.

74. Dumont, B.L., Broman, K.W., and Payseur, B.A. (2009). Variation in genomic recombination rates among heterogeneous stock mice. Genetics 182, 1345–1349. 10.1534/genetics.109.105114.

75. Dumont, B.L., and Payseur, B.A. (2011). Genetic analysis of genome-scale recombination rate evolution in house mice. PLoS Genet. 7, e1002116. 10.1371/journal.pgen.1002116.

76. Nishant, K.T., Chen, C., Shinohara, M., Shinohara, A., and Alani, E. (2010). Genetic analysis of baker’s yeast Msh4-Msh5 reveals a threshold crossover level for meiotic viability. PLoS Genet. 6. 10.1371/journal.pgen.1001083.

77. Shodhan, A., Lukaszewicz, A., Novatchkova, M., and Loidl, J. (2014). Msh4 and Msh5 function in SC-independent chiasma formation during the streamlined meiosis of Tetrahymena. Genetics 198, 983–993. 10.1534/genetics.114.169698.

78. Kelly, K.O., Dernburg, A.F., Stanfield, G.M., and Villeneuve, A.M. (2000). *Caenorhabditis elegans* msh-5 is required for both normal and radiation-induced meiotic crossing over but not for completion of meiosis. Genetics 156, 617–630. 10.1093/genetics/156.2.617.

79. Hollingsworth, N.M., Ponte, L., and Halsey, C. (1995). MSH5, a novel MutS homolog, facilitates meiotic reciprocal recombination between homologs in Saccharomyces cerevisiae but not mismatch repair. Genes Dev. 9, 1728–1739. 10.1101/gad.9.14.1728.

80. He, W., Rao, H.B.D.P., Tang, S., Bhagwat, N., Kulkarni, D.S., Ma, Y., Chang, M.A.W., Hall, C., Bragg, J.W., Manasca, H.S., et al. (2020). Regulated proteolysis of mutsγ controls meiotic crossing over. Mol. Cell 78, 168–183.e5. 10.1016/j.molcel.2020.02.001.

81. Lam, I., and Keeney, S. (2014). Mechanism and regulation of meiotic recombination initiation. Cold Spring Harb. Perspect. Biol. 7, a016634. 10.1101/cshperspect.a016634.

82. Sanchez, A., Adam, C., Rauh, F., Duroc, Y., Ranjha, L., Lombard, B., Mu, X., Wintrebert, M., Loew, D., Guarné, A., et al. (2020). Exo1 recruits Cdc5 polo kinase to MutLγ to ensure efficient meiotic crossover formation. Proc Natl Acad Sci USA 117, 30577–30588. 10.1073/pnas.2013012117.

83. Silva, N., Adamo, A., Santonicola, P., Martinez-Perez, E., and La Volpe, A. (2013). Pro-crossover factors regulate damage-dependent apoptosis in the Caenorhabditis elegans germ line. Cell Death Differ. 20, 1209–1218. 10.1038/cdd.2013.68.

84. He, W., Verhees, G.F., Bhagwat, N., Yang, Y., Kulkarni, D.S., Lombardo, Z., Lahiri, S., Roy, P., Zhuo, J., Dang, B., et al. (2021). SUMO fosters assembly and functionality of the MutSγ complex to facilitate meiotic crossing over. Dev. Cell 56, 2073–2088.e3. 10.1016/j.devcel.2021.06.012.

85. Pittman, D.L., Cobb, J., Schimenti, K.J., Wilson, L.A., Cooper, D.M., Brignull, E., Handel, M.A., and Schimenti, J.C. (1998). Meiotic prophase arrest with failure of chromosome synapsis in mice deficient for Dmc1, a germline-specific RecA homolog. Mol. Cell 1, 697–705. 10.1016/s1097-2765(00)80069-6.

86. Yoshida, K., Kondoh, G., Matsuda, Y., Habu, T., Nishimune, Y., and Morita, T. (1998). The mouse RecA-like gene Dmc1 is required for homologous chromosome synapsis during meiosis. Mol. Cell 1, 707–718. 10.1016/s1097-2765(00)80070-2.

87. Shin, Y.-H., McGuire, M.M., and Rajkovic, A. (2013). Mouse HORMAD1 is a meiosis i checkpoint protein that modulates DNA double-strand break repair during female meiosis. Biol. Reprod. 89, 29. 10.1095/biolreprod.112.106773.

88. Rinaldi, V.D., Bolcun-Filas, E., Kogo, H., Kurahashi, H., and Schimenti, J.C. (2017). The DNA damage checkpoint eliminates mouse oocytes with chromosome synapsis failure. Mol. Cell 67, 1026–1036.e2. 10.1016/j.molcel.2017.07.027.

89. Bolcun-Filas, E., Rinaldi, V.D., White, M.E., and Schimenti, J.C. (2014). Reversal of female infertility by Chk2 ablation reveals the oocyte DNA damage checkpoint pathway. Science 343, 533–536. 10.1126/science.1247671.

90. Swindells, E.O.K., Alesi, L.R., Stringer, J.M., and Hutt, K.J. (2026). Guidelines for quantifying ovarian follicles: every follicle counts. Reproduction 171. 10.1093/reprod/xaag002.

91. Peters, A.H., Plug, A.W., van Vugt, M.J., and de Boer, P. (1997). A drying-down technique for the spreading of mammalian meiocytes from the male and female germline. Chromosome Res. 5, 66–68. 10.1023/a:1018445520117.

92. Holloway, J.K., Morelli, M.A., Borst, P.L., and Cohen, P.E. (2010). Mammalian BLM helicase is critical for integrating multiple pathways of meiotic recombination. J. Cell Biol. 188, 779–789. 10.1083/jcb.200909048.

93. Horan, T.S.A. (2025). Preparation of Meiotic Chromosome Spreads from Mouse Oocytes for Assessment of Synapsis and Recombination. J. Vis. Exp. 10.3791/68749.

94. Sun, X., and Cohen, P.E. (2013). Studying recombination in mouse oocytes. Methods Mol. Biol. 957, 1–18. 10.1007/978-1-62703-191-2_1.

95. urner, J.M.A., Aprelikova, O., Xu, X., Wang, R., Kim, S., Chandramouli, G.V.R., Barrett, J.C., Burgoyne, P.S., and Deng, C.-X. (2004). BRCA1, histone H2AX phosphorylation, and male meiotic sex chromosome inactivation. Curr. Biol. 14, 2135–2142. 10.1016/j.cub.2004.11.032.

