## Supplementary figures and images for "A mammalian-specific domain of MSH5 drives the transition from crossover licensing to designation during meiotic prophase I"

### Figure S1

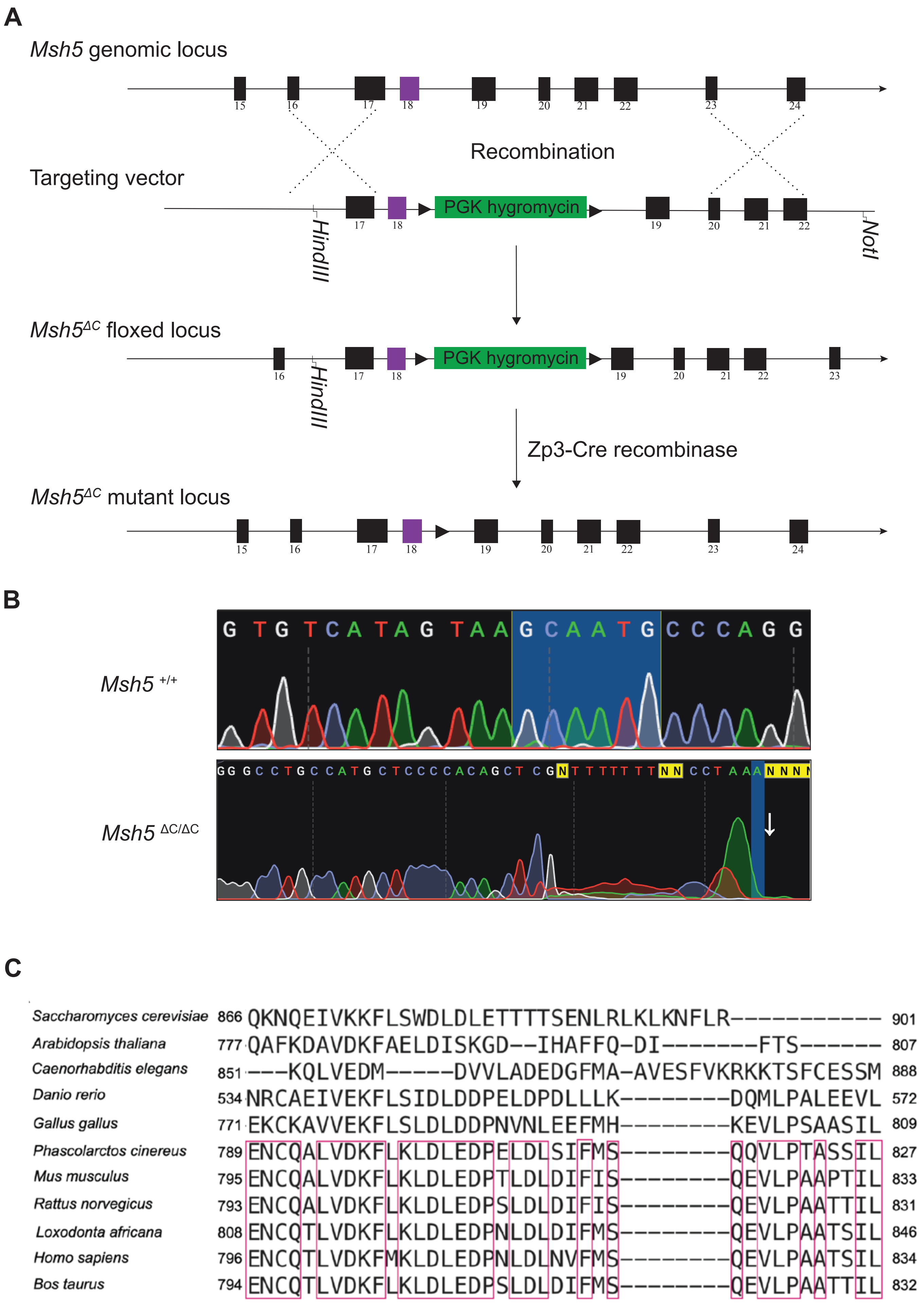

### Figure S2

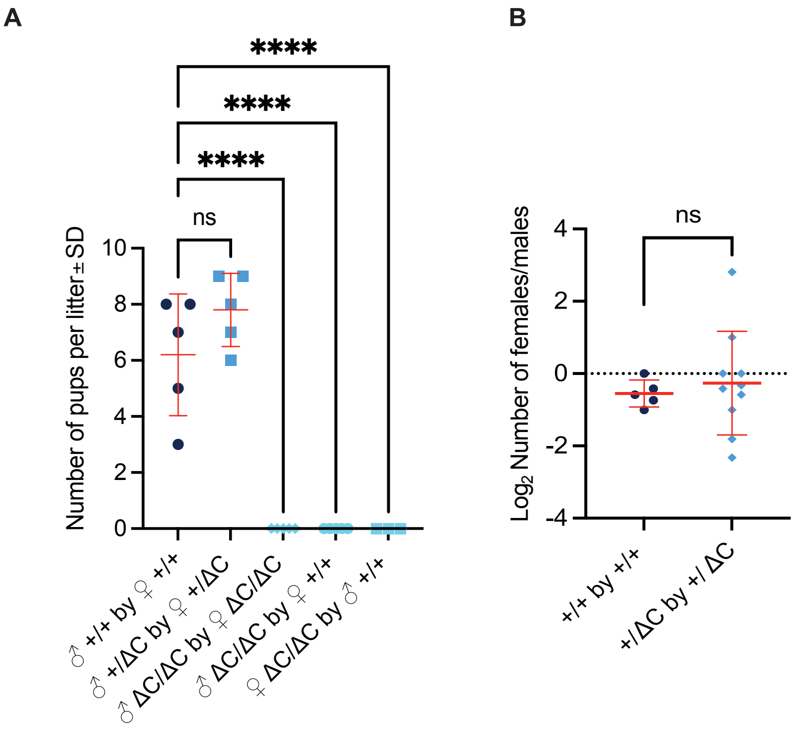

### Figure S3

**Carr *et al*.**
**FIGURE S3**

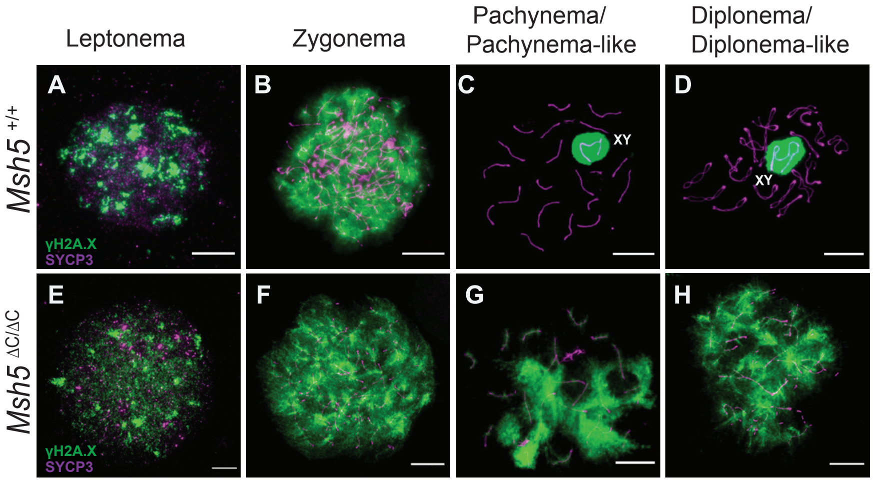

### Figure S4

Carr *et al.*
**FIGURE S4**

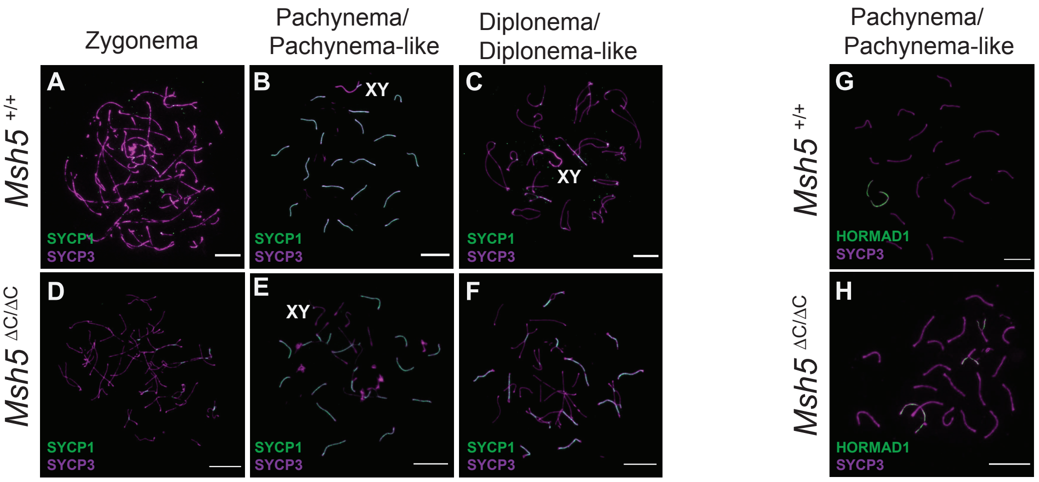

### Figure S6

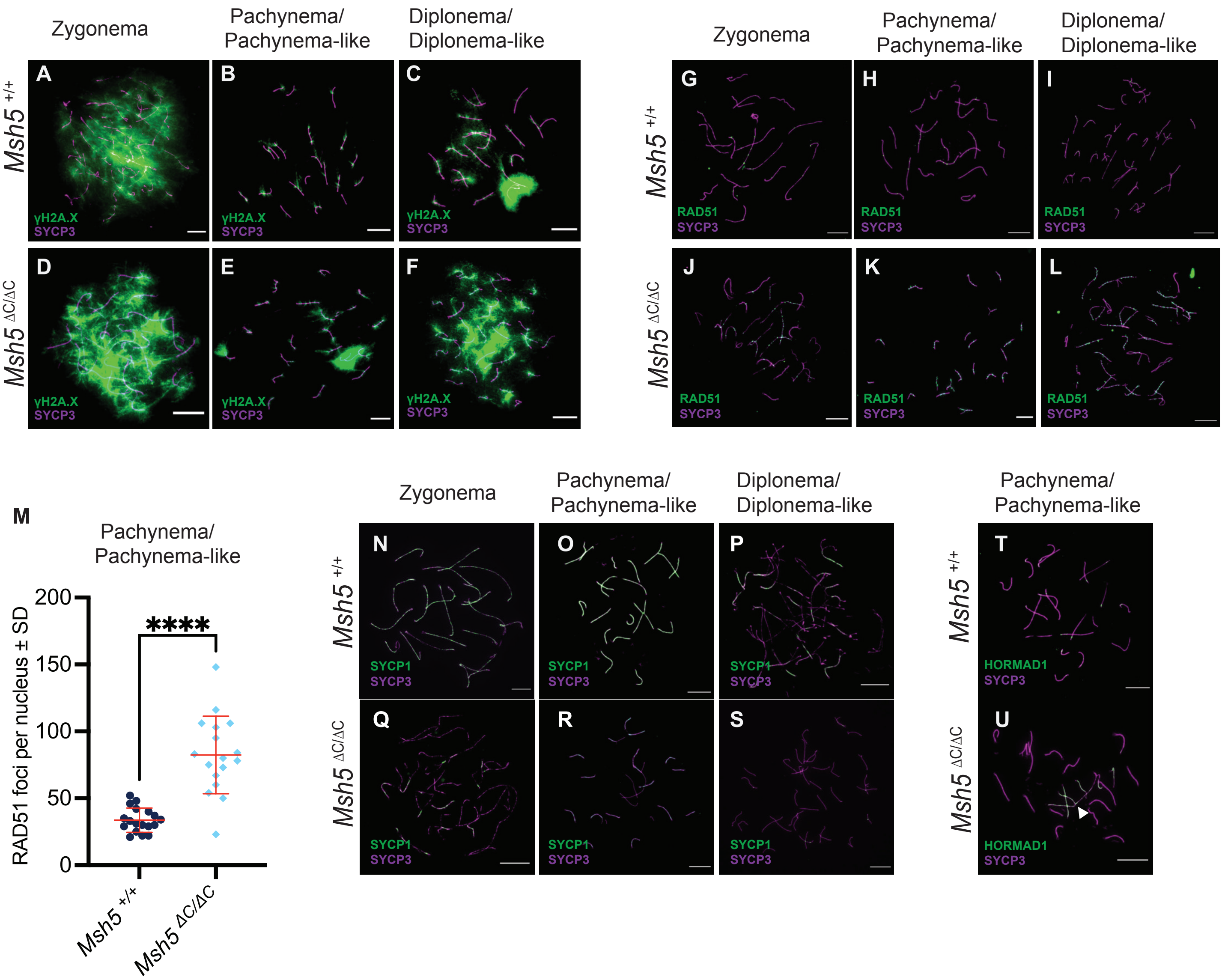

### Figure S6

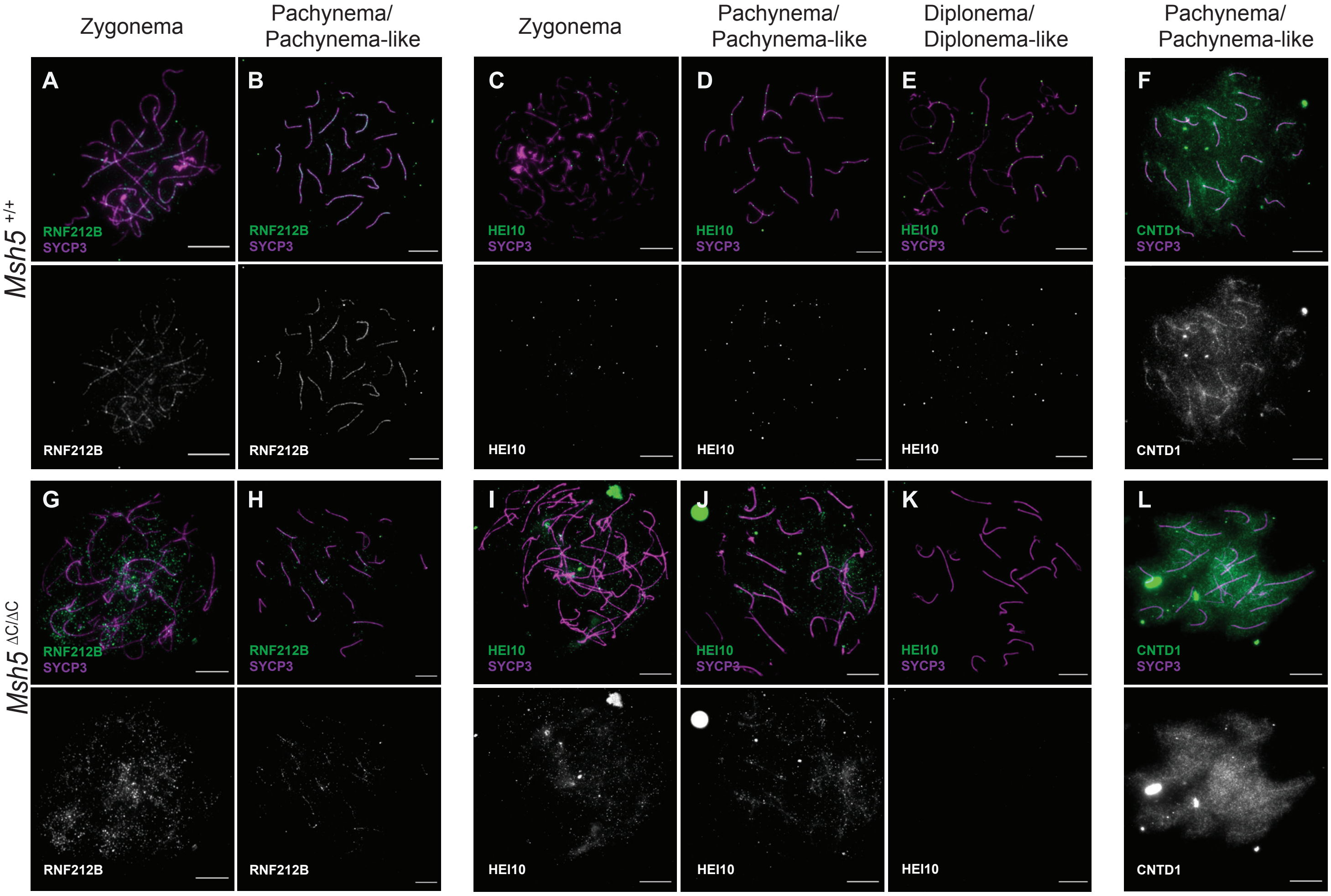

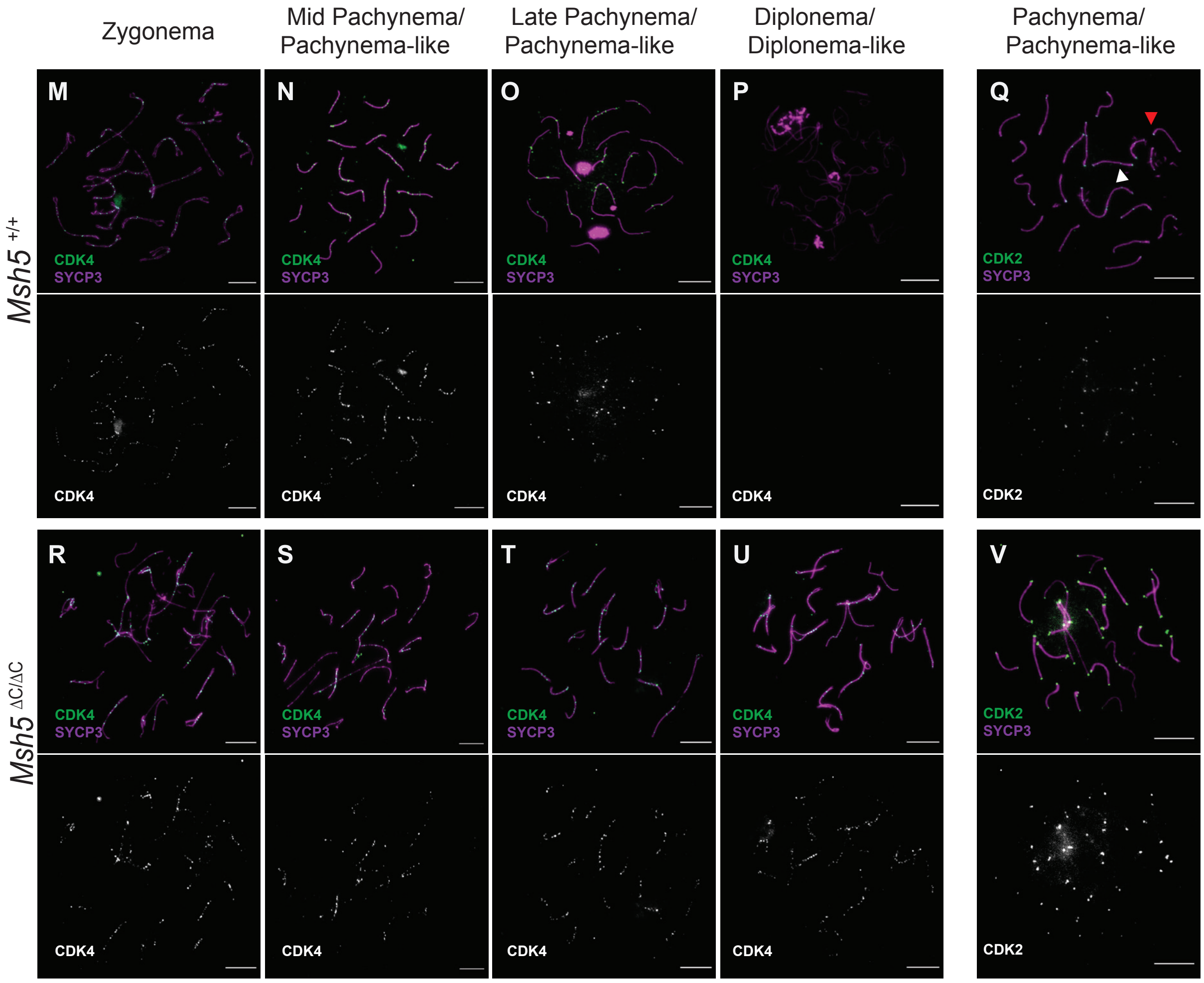
